# Evaluation of haplotype callers for next-generation sequencing of viruses

**DOI:** 10.1101/828350

**Authors:** Anton Eliseev, Keylie M. Gibson, Pavel Avdeyev, Dmitry Novik, Matthew L. Bendall, Marcos Pérez-Losada, Nikita Alexeev, Keith A. Crandall

**Author notes:** Co-First Author (Anton Eliseev, Keylie M. Gibson). Co-Second Author (Pavel Avdeyev, Dmitry Novik). Joint Senior Authors (Nikita Alexeev, Keith A. Crandall). Corresponding author at: Computational Biology Institute, Milken Institute School of Public Health, George Washington University, Washington, DC, USA, Email address (K.M. Gibson).

## Abstract

Currently, the standard practice for assembling next-generation sequencing (NGS) reads of viral genomes is to summarize thousands of individual short reads into a single consensus sequence, thus confounding useful intra-host diversity information for molecular phylodynamic inference. It is hypothesized that a few viral strains may dominate the intra-host genetic diversity with a variety of lower frequency strains comprising the rest of the population. Several software tools currently exist to convert NGS sequence variants into haplotypes. However, previous studies suggest that current approaches of haplotype reconstruction greatly underestimate intra-host diversity. Here, we tested twelve NGS haplotype reconstruction methods using viral populations simulated under realistic evolutionary dynamics. Parameters for the simulated data spanned known fast evolving viruses (e.g., HIV-1) diversity estimates to test the limits of the haplotype reconstruction methods and ensured coverage of predicted intra-host viral diversity levels. Using those parameters, we simulated HIV-1 viral populations of 216-1,185 haplotypes per host at a frequency <7%. All twelve investigated haplotype callers showed variable performance and produced drastically different results that were mainly driven by differences in mutation rate and, to a lesser extent, in effective population size. Most methods were able to accurately reconstruct haplotypes when genetic diversity was low. However, under higher levels of diversity (e.g., those seen intra-host HIV-1 infections), haplotype reconstruction accuracy was highly variable and, on average, poor. High diversity levels led to severe underestimation of, with a few tools greatly overestimating, the true number of haplotypes. PredictHaplo and PEHaplo produced estimates close to the true number of haplotypes, although their haplotype reconstruction accuracy was worse than that of the other ten tools. We conclude that haplotype reconstruction from NGS short reads is unreliable due to high genetic diversity of fast-evolving viruses. Local haplotype reconstruction of longer reads to phase variants may provide a more reliable estimation of viral variants within a population.

**Highlights:**

- Haplotype callers for NGS data vary greatly in their performance.
- Haplotype callers performance is mainly determined by mutation rate.
- Haplotype callers performance is less sensitive to effective population size.
- Most haplotype callers perform well with low diversity and poorly with high diversity.
- PredictHaplo performs best if genetic diversity is in the range of HIV diversity.

## 1. Introduction

Next-generation sequencing (NGS) technologies provide novel opportunities to study the evolution of many viruses that impose health issues among humans, such as human immunodeficiency virus (HIV), hepatitis C virus (HCV), human papillomavirus (HPV), and influenza. Such sequencing platforms allow an in-depth characterization of the genetic diversity in a heterogeneous intra-host viral population by sequencing many viral strains directly. Illumina and 454/Roche offered the first round of next-generation sequencing machines, which gradually replaced Sanger sequencing for viral studies. These platforms are able to generate a sufficiently high coverage of the genome, which allows one to detect mutations present in less abundant strains. However, the large number of short reads with a relatively high error rate produced during sequencing poses computational and statistical challenges for reconstructing full-length strain sequences and estimating their frequency. In particular, since abundance rates can be comparable or lower than sequencing error rates, high sequence error rates (≤0.1% for Illumina reads) can interfere with the detection of true mutations that are present at low frequencies. Moreover, short reads length (25 – 400 bp) need to be assembled into an unknown number of contigs. Ultimately, the goal of assembly is to produce contigs that can cover the entire targeted gene region (i.e., targeted amplicon sequencing) or that can be scaffolded together to cover the length of a full genome (i.e., shotgun sequencing). Finally, the large number of sequencing reads (25 – 300 million) requires the development of algorithms capable of processing this large amount of data. The amount of data generated by a single NGS run (1 GB to 1 TB) can be up to a million times greater than that generated by a single Sanger sequencing run (1 MB of data).

Several computational tools have been developed over the last decade to address the challenge of defining sequence variants (haplotypes – sometimes erroneously referred to as ‘quasispecies’; see Holmes, 2010) from NGS data (Beerenwinkel and Zagordi, 2011; Di Giallonardo et al., 2014; Pandit and de Boer, 2014; Posada-Cespedes et al., 2016; Schirmer et al., 2014). Different software has been tailored to various sequencing platforms and experimental designs. It is important to note that 454/Roche sequencing reads were the main input data for developers of viral variant assemblers until 2013. This was because 454/Roche was the first widely-used NGS platform and generated longer reads than all other Illumina platforms at the time (Beerenwinkel and Zagordi, 2011; Schirmer et al., 2014). A number of computational methods were proposed for handling the 454/Roche reads, including PredictHaplo (Prabhakaran et al., 2014), ViSpA (Astrovskaya et al., 2011), QuRe (Prosperi and Salemi, 2012), QuasiRecomb (Topfer et al., 2013), VirA (Skums et al., 2013), BIOA (Mancuso et al., 2011), Mutant-Bin (Prabhakara et al., 2013), V-Phaser + V-Profiler^1^ (Henn et al., 2012; Macalalad et al., 2012), and ShoRAH (Zagordi et al., 2010). Some of these methods were empirically validated using HIV and HCV data sets with the methods showing little success in estimating reliable sequence variants from NGS data (Prosperi et al., 2013). Later, thanks to the better cost-effectiveness and higher coverage offered by the Illumina sequencing platforms, the main focus migrated towards Illumina technology and has become dominant for developers of viral sequence variant assemblers since then (Posada-cespedes et al., 2016). Following this paradigm shift, several methods such as PredictHaplo, V-Phaser (Yang et al., 2013), and QuasiRecomb were extended to handle Illumina reads, and a number of tools, including VGA (Mangul et al., 2014), HaploClique (Töpfer et al., 2014), QColors (Huang et al., 2011), QSdpR (Barik et al., 2018), and ViQuas (Jayasundara et al., 2015), were developed specifically to handle Illumina reads.

Currently, all state-of-the-art methods for viral variant reconstruction are designed to assemble contigs from Illumina reads and can be divided into two main categories based on their dependency on a reference genome: *reference-based* assemblers and *de novo* assemblers (Fig. 1). In the former category, sequencing reads are aligned to a reference genome and information about the reads positioning and orientation with respect to a reference genome is obtained (Fig. 1 c_1_). This information is further used to reconstruct haplotypes (Fig. 1 c_2_, c_3_, d_1_, d_2_). *De novo* assemblers, however, do not rely on reference genomes, and haplotype sequences are usually reconstructed directly from the reads (Fig. 1 c_4_, d_3_). *De novo* assembly often requires more computational resources, but *reference-based* assembly requires a closely related genome, which is not always available in high quality. Each strategy has advantages and disadvantages and have been implemented in several software programs, but the performance of these assembly tools has not been comprehensively examined yet. In this study, we simulated realistic, coalescent based intra-host viral diversity with diversity measurements encompassing known variation from fast-evolving viruses such as HIV-1 for empirical grounding. We then used these simulated populations to assess the performance and accuracy of three *de novo* and nine *reference-based* sequence variant reconstruction tools or haplotype callers.

**Figure 1.**
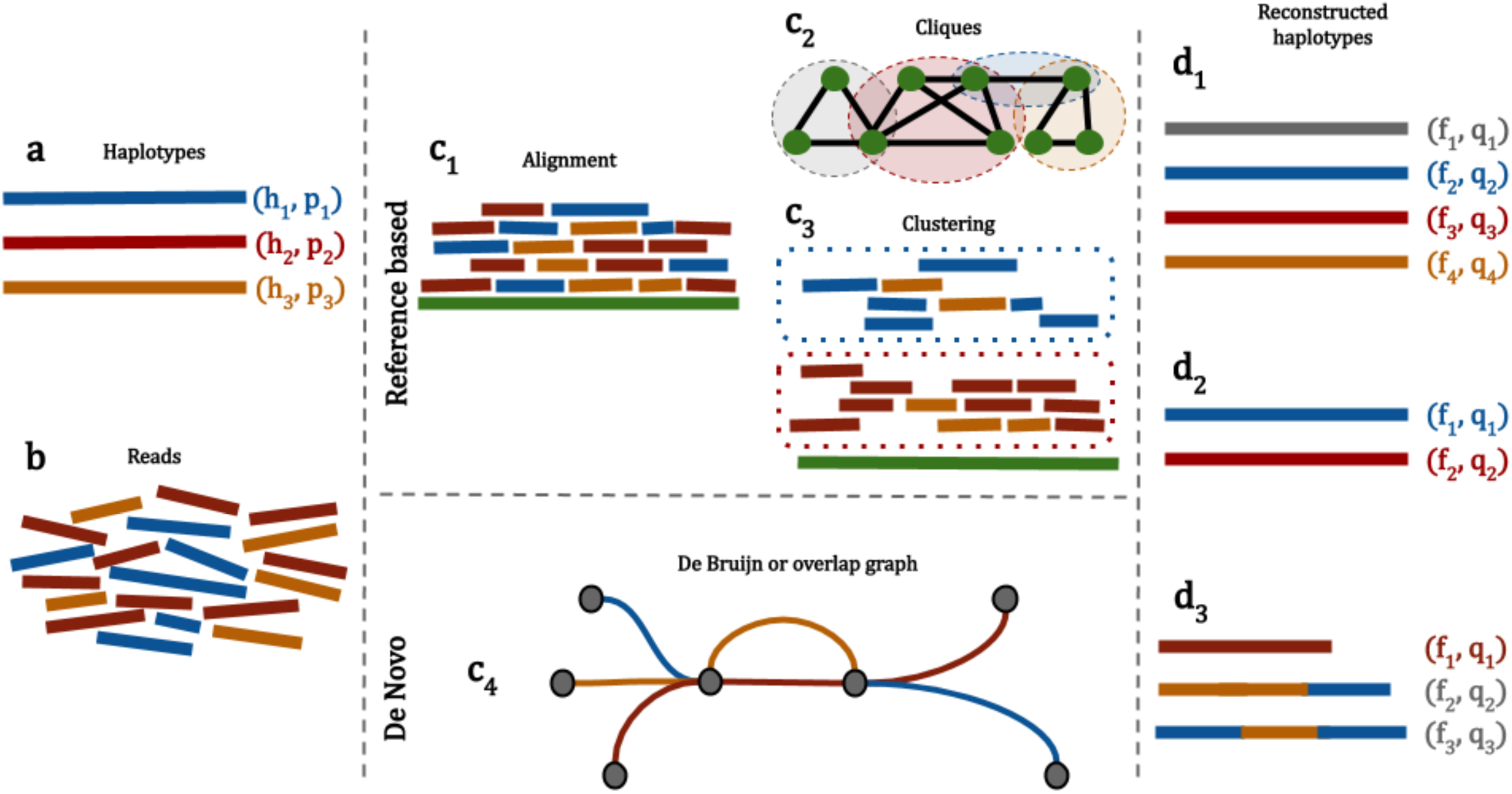
Schematic diagram representing the process of reconstructing haplotypes from next-generation sequencing reads by reference-based and *de novo* methods. (a) A hypothetical virus population consisting of three haplotypes is sequenced by NGS techniques. (b) Reads originating from different haplotypes are identified by distinct colors in the diagram. (c_1_) After sequencing, reads are aligned against reference genome (green) as a first step in all reference-based methods. (c_2_) Read alignment is used for building a graph and candidate haplotypes are reconstructed as maximal cliques in the graph. (c_3_) Read alignment is used for dividing reads into clusters and candidate haplotypes are reconstructed by concatenation of all reads from clusters. (c_4_) Alternatively, after sequencing, reads are *de novo* assembled using De Bruijn or overlap graphs and candidate haplotypes are reconstructed as paths by analyzing the graph structure. (d_1_) A method based on clique detections overestimates the number of reconstructed haplotypes with relative frequencies. (d_2_) A method based on clustering procedure underestimates the number of reconstructed haplotypes with relative frequencies. (d_3_) A *de novo* method reconstructs the correct number of haplotypes with frequencies, but one inferred haplotype is smaller than the true haplotype and the other two haplotypes are admixtures of the original haplotypes.

## 2. Material and Methods

### 2.1 Viral Sequence Variant Estimators

All *de novo* viral variant reconstruction methods can be further divided into two subcategories: *consensus* and *strain-specific* assemblers. The main goal of consensus-based tools is, generally speaking, to construct a suitable consensus reference genome that may be further used as a template for more fine-grained studies. VICUNA (Yang et al., 2012) and IVA (Hunt et al., 2015) represent this subcategory of methods. VICUNA is the most popular software among them, as it generates full-length consensus and detects polymorphisms. VICUNA merges NGS reads into contigs, and those into a bigger contig, by calculating “good” prefix-suffix overlap between sequences. During this process, contigs are also clustered and validated to reach a better quality of consensus. IVA follows the same approach with only one difference, the tool starts from k-mers that are sorted in decreasing order with respect to their abundance and then extends sequences into a bigger sequence by using reads that have perfect overlap with initial sequences. VICUNA also has an additional option for contig merging if a reference genome exists.

Contrary to *de novo* consensus approaches, *de novo* strain-specific assemblers aim to reconstruct sequences at the strain resolution level (Table 1). It is worth mentioning that the *de novo* viral variant reconstruction problem is quite similar to the assembly effort of multiple genomes in microbial communities at once using shotgun metagenomic reads (e.g., Bishara et al., 2018; Scholz et al., 2016). The arising challenges in the microbial community genome assembly are addressed by metagenome assemblers. Thus, at first glance, applying metagenome assemblers to *de novo* viral variant reconstruction seems very promising. However, SPAdes is the only assembler that was able to identify haplotypes in the case of sufficiently abundant strains (Jasmijn A Baaijens et al., 2017; Bankevich et al., 2012; Nurk et al., 2017). Therefore, the development of specific assemblers for viral sequence variants is required. Currently, there exist three *de novo* strain-specific assemblers, namely MLEHaplo (Malhotra et al., 2015), SAVAGE (Jasmijn A Baaijens et al., 2017), and PEHaplo (Chen et al., 2018) (Table 1). MLEHaplo was the first assembler that truly applied *de novo* viral sequence variant assembly at the strain resolution level. MLEHaplo performs k-mer counting and then filters erroneous k-mers using raw reads and a specified threshold value. Afterwards, the tool builds a De Bruijn graph (see Compeau et al., 2011) based on the set of k-mers obtained in the previous round (Fig. 1 c_4_). On the next step, MLEHaplo recovers paths from the De Bruijn graph that may correspond to haplotypes. Finally, the tool chooses correct haplotypes and estimates their frequencies using the maximum likelihood method. PEHaplo follows the same workflow as MLEHaplo. However, PEHaplo constructs an overlap graph instead of creating a De Bruijn graph during the initial steps^2^ (Fig. 1 c_4_). PEHaplo also has a more careful path finding algorithm based on paired-end connection information. SAVAGE uses overlap graphs as a key data structure, but the pipeline is different from those in PEHaplo and MLEHaplo. After constructing an overlap graph (Fig. 1 c_4_), SAVAGE joins overlapped read pairs. At the next step, SAVAGE iteratively merges reads into contigs and contigs into scaffolds using clique enumeration and contig formation. Finally, the tool uses Kallisto (Bray et al., 2016) to estimate frequencies of the resulting haplotypes

**Table 1.** *De novo* and *reference-based* viral haplotype callers compared in this study.

| Software Tool | Published Year | Programming Language | Frequencies | Haplotypes | Type |
| --- | --- | --- | --- | --- | --- |
| MLEHaplo | 2015 | Perl | + | + | *de novo* |
| Savage | 2017 | Python 3 | - | - | *de novo* |
| PEHaplo | 2018 | Python 2.7 | + | - | *de novo* |
| ShoRAH | 2011 | C++ | + | + | *reference-based* |
| QuRe | 2012 | Java 6 | + | + | *reference-based* |
| QuasiRecomb | 2013 | Java 7 | + | + | *reference-based* |
| PredictHaplo | 2014 | C++ | + | + | *reference-based* |
| HaploClique | 2014 | C++ | + | + | *reference-based* |
| ViQuas | 2015 | R, Perl | + | + | *reference-based* |
| aBayesQR | 2017 | C++ | + | + | *reference-based* |
| RegressHaplo | 2017 | R | + | + | *reference-based* |
| CliqueSNV | 2018 | Java 6 | + | + | *reference-based* |

If a haplotype caller produces haplotypes as an output, it is given a plus sign. If a haplotype caller reports corresponding frequencies for the sequences produced, it is given a plus sign. Savage and PEHaplo claim they produce contigs not haplotypes, which is why we did not deem that they produce haplotypes.

While the final sequences produced by MLEHaplo, PEHaplo, and SAVAGE are strain-specific, the obtained sequences, in general, do not represent full-length haplotypes (Fig. 1 d_3_). Recently, Virus-VG and VG-flow have been developed for completing strain-specific assemblies produced by the aforementioned *de novo* strain-specific assemblers (Baaijens et al., 2018, 2019). Virus-VG and VG-flow try to convert strain-specific contigs into full-length haplotypes taking into account their abundances. The difference between Virus-VG and VG-flow is that the former uses a brute-force exact approach, while the latter utilizes a heuristic algorithm. Therefore, VG-flow is faster than Virus-VG, but less accurate. The main goal for both tools is to find and select maximum-length paths in a variation graph.

The main advantage of *reference-based viral* variant reconstruction methods prior to *de novo* haplotype assemblers is the potential ability to reconstruct full-length haplotypes (Fig. 1 d_1_, d_2_). However, it was shown in several studies (Jasmijn A Baaijens et al., 2017; Mangul et al., 2014) that the reference genome may bias the reconstruction of haplotypes. An additional disadvantage of using a reference-based tool is the potential lack of a high-quality reference genome of a virus population. In this case, the required reference genome can be potentially assembled from sequencing reads by first using *de novo* consensus assembly tools. Nevertheless, the reference genome is often available for common pathogenic viruses, such as HIV, HCV, polyomavirus or influenza.

Currently there are nine commonly used state-of-the-art reference-based tools (Table 1). All these tools claim to be global haplotype inference methods, i.e., able to infer the sequences and frequencies of the underlying viral strains over a longer region than the average read length. ShoRAH (Short Read Assembly into Haplotypes) is, historically, the first publicly available software (Zagordi et al., 2011). ShoRAH uses a probabilistic clustering algorithm for short haplotype sequence reconstruction (Fig. 1 c_1_, c_3_, d_2_). Then, it computes a minimal set of haplotypes using the principle of parsimony that provides the best explanation for the given a set of error corrected sequencing reads (Eriksson et al., 2008). The tool then uses an expectation minimization algorithm for haplotype frequency estimation.

The next important milestone in the reference-based viral variant reconstruction tool development was the release of QuRe (Prosperi and Salemi, 2012). QuRe uses the combinatorial method proposed in Prosperi and Salemi (2012) for inferring genetic variants in local windows that do not exceed read lengths. After that, the obtained genetic variants are clustered by a probabilistic algorithm (Zagordi et al., 2010) (Fig. 1 c_1_, c_3_, d_2_). Finally, haplotypes with their frequencies are obtained by utilizing a genome reference and clustered variants. Later, the same probabilistic clustering and combinatorial algorithms were used for developing the reference-assisted assembly pipeline ViQuas (Jayasundara et al., 2015). The main difference between QuRe and ViQuas is that the latter tool assembles reads into contigs using the SSAKE assembler (Warren et al., 2007) and then iteratively extends obtained contigs by connecting overlapping pairs without using any sequence information from the reference.

The next developed software was PredictHaplo (Prabhakaran et al., 2014), HaploClique (Töpfer et al., 2014), and QuasiRecomb (Topfer et al., 2013). All these tools have special features in comparison to the previous generation tools. For example, PredictHaplo was specifically designed for identifying haplotypes in an HIV-1 population. HaploClique allows for detection of point mutations, large insertions and deletions. QuasiRecomb, on the other hand, incorporates the existence of recombination events into the estimated viral evolution. PredictHaplo, HaploClique, and QuasiRecomb are based on different approaches and their applications to the viral variant reconstruction problem were novel at the time. PredictHaplo reformulates the original problem in terms of a non-standard clustering problem, where reads are points in some metric space and haplotypes are clusters (Fig. 1 c_1_, c_3_, d_2_). To take into account an unknown number of variants, the stochastic Dirichlet process and the infinite mixture model were used (Prabhakaran et al., 2010). HaploClique uses the insert size distribution and an iterative enumeration of maximal cliques in a graph to reconstruct super-reads that may represent haplotypes (Fig. 1 c_1_, c_2_, d_1_). Due to the computational complexity of maximal clique enumeration, this tool requires excessive computational resources on data sets with coverage >1,000x. HaploClique provided inspiration for the development of SAVAGE. Finally, QuasiRecomb utilizes data parameters of a hidden Markov model for estimating point mutations and recombination events (David Posada et al., 2002). These parameters allow estimation of the probability of each possible haplotype with respect to the observed read data.

The latest releases of reference-based methods for viral sequence variant reconstruction are aBayesQR (Ahn and Vikalo, 2017), CliqueSNV (Knyazev et al., 2018), and RegressHaplo (Leviyang et al., 2017). CliqueSNV extends the previous approach used in the 2SNV tool (Artyomenko et al., 2017). CliqueSNV constructs a graph based on linkage information between single nucleotide variations and then identifies true viral variants by merging cliques in that graph (Fig. 1 c_1_, c_2_, d_1_). RegressHaplo, in turn, is based on a regression-based approach specifically designed low diversity and convergent evolutions. This tool implements penalized regression to assess the haplotype interactions that belong to different unlinked regions. aBayesQR employs a maximum-likelihood approach to infer viral sequences. The search of most likely viral sequence is conducted on long contigs, which enables identification of closely related haplotypes in a population and provides computational tractability of the Bayesian method. It should be noted that aBayesQR is designed for reconstructing viral haplotypes that are near genetically identical.

Each haplotype reconstruction tool in Table 1 was run on the Colonial One high performance computing cluster at The George Washington University. We used 64 standard CPU nodes featuring dual Intel Xeon E5-2670 2.6GHz 8-core processors with a RAM capacity of 128GB. A single node with a 48-hour time limit was allocated for each run.

### 2.2 Simulation Data Description

Previous benchmarking of viral haplotype reconstruction programs (Pandit and de Boer, 2014; Prosperi et al., 2013; Schirmer et al., 2014) used simulation scenarios that are useful from a mathematical perspective but do not necessarily reflect viral evolution and epidemiology. For example, PredictHaplo artificially mutated ten haplotypes from a single HIV-1 reference genome at varying proportions (Prabhakaran et al., 2014); HaploClique used an in-house mixture of known HIV-1 strains (number of specific strains unknown) (Töpfer et al., 2014); and SAVAGE simulated their data based on Illumina MiSeq sequencing results from an in-house mixture of five unique strains of HIV-1 subtype B with varying relative abundances (see supplemental methods in (Jasmijn A Baaijens et al., 2017)). In those studies, often the pairwise divergence between the strains used to represent “real” HIV-1 haplotype diversity was either unreported (Prabhakaran et al., 2014) or ranged between 0.05% and 10% (Jasmijn A Baaijens et al., 2017; Töpfer et al., 2014). But realistic intra-host HIV-1 diversity is substantially lower with pairwise divergences ranging between 0.02% and 2%, while inter-host pairwise comparisons of the same viral subtypes can exceed 5% (Kearney et al., 2009; Maldarelli et al., 2013). Furthermore, unless the HIV-1 viral population in an individual was the product of a dual infection (see (van der Kuyl and Cornelissen, 2007) for review of dual infections), these benchmarking methods do not accurately represent the evolution of the virus, where the HIV viral population originated from an infection of one strain. All of these studies conditioned their simulations on HIV-1 data sets, but we also want to explore the general utility across a broader parameter space that encompasses more fast-evolving viral populations.

In our simulations, we used parameters and settings under the coalescent theory (Kingman, 2000, 1982; Rodrigo and Felsenstein, 1999; Rosenberg and Nordborg, 2002) to more accurately reflect viral intra-host diversity and evolution as seen in empirical studies (see (Crandall and Templeton, 1993)). We simulated viral intra-host evolutionary histories and the constituent haplotype sequences (tips) using the coalescent simulator CoalEvol v. 7.3.5 (Arenas and Posada, 2014). We set the mutation rate (*μ*) between 1e-3 and 5e-8 per-site to span past known viral mutation rates to test the limits of the reconstruction algorithms and number of haplotypes present using the human genome mutation rate as an upper limit and other retroviruses’ mutation rates as a lower limit. These parameters encapsulated the empirical mutation rate of 2.5e-5 and 3.4e-5 estimated by Neher and Leitner (2010) and Maldarelli et al. (2013), respectively for HIV-1; HCV with an estimated mutation rate between 2.5e-5 and 1.2e-4 (Echeverría et al., 2015; Ribeiro et al., 2012; Sanjuán et al., 2010); HTLV-1 with an estimated mutation rate between 3.44e-7 and 7e−6 (Mansky, 2000; Nobre et al., 2018); and influenza with an estimated mutation rate of 3e−5 to 4e−5 (McCrone, 2018; McCrone et al., 2018). Although Neher and Leitner (2010) reported that the HIV-1 virus recombines at a rate of 1.4±0.6e−5, we chose to not include recombination in the simulated evolution histories because some of the compared haplotype reconstruction programs do not include recombination events in their reconstruction process. Other parameters that were fixed in the CoalEvol config file included: i) nucleotide frequencies (A=0.37, C=0.16, G=0.23, and T=0.25); ii) the transition/transversion rate (ti/tv = 2.5), as estimated among host diversity from Crandall et al. (1999); and iii) rate heterogeneity among sites (Γ = 0.95) and invariable site rate (I = 0.4) (Posada and Crandall, 2001), which are unique to HIV-1.

Recombination occurs frequently in natural HIV-1 populations (Crandall, 1999; Neher and Leitner, 2010; D Posada et al., 2002) but we chose not to model recombination in our simulations. First, many of the haplotype programs do not account for recombination. Second, we assume that approaches that fail on a simplified model without recombination will not perform well on a more complex that includes recombination.

We chose to use HIV-1 as an empirical viral strain to assess the capabilities of the haplotype reconstruction tools given that most developers validated their programs on this virus and genetic diversity values for this virus are well established. HIV-1 genetic diversity (Watterson’s theta) for the polymerase gene (*pol*) has been estimated to fall between 0.067 and 0.09 for subtype B HIV-1 strains in the United States (Gibson et al., 2019; Pérez-Losada et al., 2017, 2010). Boltz et al. (2016) completed single genome sequencing that resulted in 677 – 1,577 sequences per sample for HIV-1, therefore, we limited our sample size to range between 100 and 2,000 with an alignment length of 1,137bp. This length was chosen because we used a section of the polymerase gene (*pol*) from the HXB2 reference sequence (GenBank accession number: K03455; (Ratner et al., 1985)) as the most recent common ancestor (MRCA) for each parameter set (HXB2 numbering: 2,253 – 3,390). It is important to note that CoalEvol is restricted to sample sizes of up to 2,000 haplotypes. Maldarelli et al. (2013) estimated the effective population size (*N*_*e*_) of intra-host diversity to be between 1,000 and 10,000, so we varied the effective population size between 500 and 10,000. We also denoted the ploidy as diploid (Coffin, 1992). Wherever possible, we varied the parameters to be above and below estimated HIV-1 estimates to ensure we adequately represented viral intra-host diversity and to examine the limits of the haplotype reconstruction programs. Expanding our parameter set allowed us to gain insights into other viral species with different evolutionary and population characteristics. For example, the *N*_*e*_ for influenza is considerably smaller than HIV-1 at around 20-100 viral sequences (Kim and Kim, 2016; McCrone, 2018; McCrone et al., 2018), while HCV hovers around the lower end of HIV-1 with an *N*_*e*_ of 10-1,000 sequences (Bernini et al., 2011).

Since the Illumina MiSeq platform is the most popular NGS technology currently used for viral amplicon sequencing due to low cost and high throughput, we simulated sequencing reads in the FASTA output (excluding the original HXB2 sequence we deemed as the GMCRA in the coalescence simulation) of CoalEvol using the NGS read simulator ART v. MountRainier-2016-06-05 (Huang et al., 2012). ART mimics real sequencing processes, therefore, we used the built-in sequencing Illumina MiSeq platform (MSv1). We simulated error-free 150 bp paired-end reads with a read count of 100 reads, mean size of 215 bp for DNA fragments, and a standard deviation of 120 bp for DNA fragment size.

The error free output data generated for the haplotype populations with the ART read simulator was processed with HAPHPIPE, a HAplotype reconstruction and PHylodynamics PIPEline for viral NGS sequences (Bendall et al., 2019). By both not simulating recombination and starting with sequencing-error free data, we removed nuisance variables that would impact haplotype reconstruction and could not be handled by some haplotype callers. Briefly, we used HAPHPIPE and its implementation of Trimmomatic v. 0.33 (Bolger et al., 2014) to trim the starting FASTQ files from the output of ART by removing low quality reads, low quality bases, and adapter contamination. We performed *de novo* assembly on the clean reads using Trinity v. 2.5.1 (Grabher et al., 2013) and formed scaffolds with MUMMER 3+ v. 3.23 (Alnafee, 2016). With two iterative refining steps, the cleaned reads were mapped back to the scaffolds with Bowtie2 v. 2.3.4.1 (Langmead and Salzberg, 2013). The BAM file of aligned reads generated as final output by HAPHPIPE and a FASTA file containing the cleaned reads (an intermediate output by HAPHPIPE) were used as input for the haplotype reconstruction algorithms.

### 2.3 Haplotype Assembly Comparative Indices

In order to evaluate the quality of haplotype assembly provided by different tools, we used common statistical measures of precision and recall, as well as weighted normalized UniFrac distance (Lozupone and Knight, 2005), which is widely used to compare microbial communities. Our simulated data can be represented as *P* = {(*h*_*i*._, *p*_*i*._), *i* = 1, 2, …} – the ground truth haplotypes *h*_*i*._ and their associated abundances *p*_*i*._ (∑ *p*_*i*._ = 1), and *Q* = {(*f*_*i*._, *q*_*i*._), *i* = 1, 2, …} – the set of predicted haplotypes *f*_*i*_ together with their predicted abundances *q*_*i*._.

We define precision as 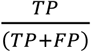 and recall as 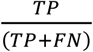. Since the length of viral sequences reconstructed by *de novo* tools may differ from actual length of ground truth haplotypes, we define *TP* (true positive) and *FP* (false positive) differently for *reference-based* and *de novo* tools. We define *FN* (false negative) as 1 − *TP*, for both assembly strategies equally.

In the case of *reference-based* methods, we define *TP* as the total frequency of those haplotypes *h* in the ground truth set *P* which have an accurate enough prediction *f* in *Q* (which means that the edit distance *d*(*h, f*) is less than some threshold *T* = *T*(*μ*)); we also define *FP* as the total frequency of those haplotypes *f* in the predicted set *Q* which do not match any haplotype *f* from the ground truth set (which means that *d*(*h, f*) ≥ *T* for all *f* ∈ *P*). We choose the threshold *T* = 12 because 12 bp is about 1% of the haplotypes’ length. We consider the haplotype *h* ∈ *P* to be reconstructed correctly if there exists a haplotype *f* ∈ *Q* such that the edit distance between them *d*(*h, f*) ≤ 12.

For *de novo* methods, we define *TP* as follows: We say that a contig *f* from *Q* is *proper* if there exists such a ground truth haplotype *h* and its substring s ∈ *h* so that the edit distance between *f* and *s* is small (less than 1% of *f* ‘s length). Then, for each ground truth haplotype *h*, we define *c*_*i*._ – the proportion of its part which is *properly* covered by contigs from *Q*. We then define *TP* as a weighted total frequency of properly predicted haplotypes 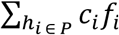. It is important to note that the definition of *TP* is a generalization of the *TP* definition for reference-based tools. Indeed, in the latter case, all the *c*_*i*_ are equal to either 0 or 1. We define *FP* as the total frequency of non-proper contigs in *Q*.

While these measures are standard and they show how good the haplotype reconstruction is, they are not very sensitive to the errors in frequency prediction. In order to address this issue, we also computed the UniFrac distance *EMD*(*P, Q*) using the EMDUniFrac algorithm (McClelland and Koslicki, 2018). The UniFrac distance takes into account both the phylogenetic structure of the haplotype set and their frequency distribution, which makes it ideal for incorporating sensitivity to errors in frequency prediction. The UniFrac EMD method makes the following steps:

- construct a tree *T* with branch length *l*_*e*_ on the set of all haplotypes *h*_*i*_ ∈ *P* and *f*_*i*_ ∈ *Q*
- for each tree branch *e* and its descendant subtree *T*_*e*_, estimates the imbalance *W*_*e*_:

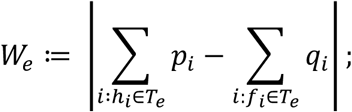
- evaluate the weighted imbalance with respect to the branch lengths

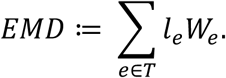

As a baseline for the UniFrac EMD comparison, we evaluate the UniFrac distance between reference or, more formally, a set of haplotypes Q containing only one haplotype – the reference at a frequency of 1.

## 3. Results and Discussion

### 3.1 True haplotypes from simulated data

All analyses were completed using the simulated dataset developed under the coalescent framework. For each mutation rate *μ* ∈{1e-8, 3e-8, 5e-8, 1e-7, 5e-7, 1e-6, 5e-6, 1e-5, 3e-5, 5e-5, 1e-4, 3e-4, 5e-4, 5e-3, 5e-8} and effective population size *N*_*e*_ = {500, 1000, 2500, 5000, 7500, 10000}, there were five simulated haplotype populations *P* = {(*h*_*i*._, *p*_*i*._), *i* = 1, 2, …} used as replicates for each parameter set. Under the coalescent model, the number of true haplotypes ranged from 1 to 1,993 with a median of 342 haplotypes for a parameter set (Fig. 2). Unlike previous attempts to represent intra-host HIV-1 diversity levels – often five haplotypes at varying abundances (J A Baaijens et al., 2017; Prabhakaran et al., 2014; Töpfer et al., 2014), our intra-host populations have 216-1,185 haplotypes per host at a frequency <7%, with a median of 525 haplotypes. Therefore, the number of haplotypes at high diversity levels may actually be even higher, but we primarily focused on the diversity levels of intra-host HIV-1 populations. Additionally, the number of haplotypes at smaller diversity levels, such as those seen in influenza, are likely to be smaller than ours.

**Figure 2.**
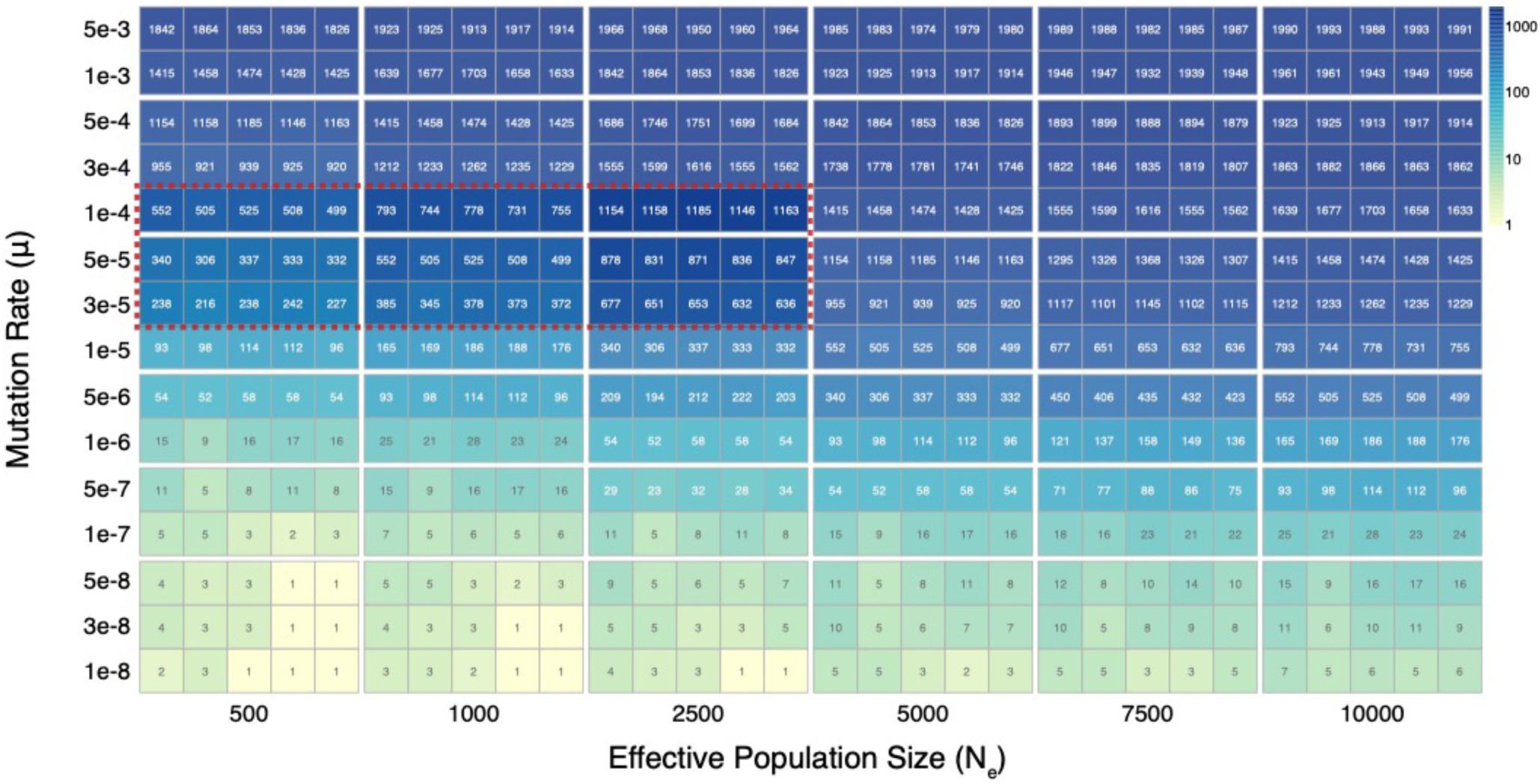
Simulated population parameters with the haplotype count in each parameter box. All five population replicates are displayed. The color darkens as the number of haplotypes increases. The red dashed box indicates expected HIV-1 mutation rates and effective population sizes.

### 3.2 Haplotype caller performance

HIV-1 intra-patient populations exhibit levels of diversity that exceed the limitations of all twelve haplotype callers we compared in this study, regardless of the assembly approach used (*de novo* or *reference-based*). However, because HCV and influenza both have lower mutation rates and effective population sizes, they may fall within the limitations of some of the compared haplotype reconstruction approaches. The haplotype callers varied drastically in their haplotype reconstruction accuracy (precision, recall, UniFrac, and number of reconstructed haplotypes), with most tools performing well with low genetic diversity and poorly with high genetic diversity. Since HIV-1 diversity is very high, all haplotype reconstruction tools seemed to have difficulties either producing output (i.e., predicted haplotypes) or reconstructing haplotypes that reflect the true haplotypes. Furthermore, haplotype reconstruction accuracy was more sensitive to the mutation rate of the virus than to its effective population size. Although, the opposite was true for PEHaplo, where *N*_*e*_ seemed to play a major role in the quality of predicted haplotypes. Fortunately, we often know, or have better *a priori* estimates for, the mutation rate of a virus than for the effective population size of an intra-host population. Furthermore, the effective population size changes over time during infection, whereas the mutation rate remains relatively constant (Maldarelli et al., 2013), unless there are pressures from antiretroviral treatment. However, as theta is estimated, effective population size and mutation rate are indeed coundfounded. Below, we discuss the current results in more detail.

MLEHaplo and ViQuas did not produce any valid results within the given time limit, whereas QuRe crashed in all analyses because of memory limitations. While HaploClique produced results within our time limit (Fig. S1), we excluded this tool from final comparisons because the length of the reconstructed viral sequences was always significantly shorter than the length of the ground truth haplotypes (Fig. S2). Such behavior is atypical among reference-based methods. Moreover, since SAVAGE can be considered as the next installment of HaploClique, it provides an additional argument for excluding HaploClique from our comparison.

In addition to the two *de novo* tools assessed (i.e., SAVAGE and PEHaplo), we also ran the VG-flow tool to complete contigs produced by those methods. We selected VG-flow over Virus-VG because VG-flow is faster and almost as accurate (Baaijens et al., 2018, 2019). Despite the claim that VG-flow improves assemblies from any *de novo* tool (Baaijens et al., 2018, 2019), we also included the output from PEHaplo in our comparison since VG-flow was tested on the SAVAGE output only in the original paper (Baaijens et al., 2018, 2019).

Datasets completed within our time limits varied across reconstruction tools; in general, datasets with higher mutation rates (*μ*) and effective population sizes (*N*_*e*_) represented challenges for almost all the tools. For example, RegressHaplo and PredictHaplo did not produce any output if both *N*_*e*_ and *μ* were high but performed well otherwise. For low values of *μ* (*μ* ≤ 1*e* − 5), all the callers except ShoRAH produced some output. aBayesQR, SAVAGE, and RegressHaplo had problems reconstructing haplotypes for datasets with low *μ* and *N*_*e*_ values.

For HIV-1 *μ* estimates (3*e* − 5 ≤ *μ* ≤ 1*e* − 4), CliqueSNV and QuasiRecomb did not produce any valid output; aBayesQR, PredictHaplo, SAVAGE, and RegressHaplo produced some outputs; and ShoRAH, and PEHaplo produced output for all the datasets. It is also important to note that only PEHaplo was able to solve all datasets within the given time limit. For all datasets, where haplotype callers performed successfully, we measured their results in terms of precision, recall, and the Unifrac distance EMD. Below we present and discuss the behavior of the *reference-based* tools and the *de novo* tools.

#### 3.2.1 Reference-based Program Performance

We evaluated results from six *reference-based* haplotype callers: aBayesQR, RegressHaplo, CliqueSNV, ShoRAH, PredictHaplo, and QuasiRecomb. Precision (Fig. 3) and recall (Fig. 4) values were calculated for each tool. The quality of obtained results did not seem to depend much on the effective population size (*N*_*e*_). This is a positive finding, as determining the effective population size for intra-host viral infections is often difficult and can vary between studies. All the tools, except ShoRAH, performed very well (i.e., both precision and recall are close to one) if the mutation rate was relatively small (*μ* ≤ 1e − 5), which is an estimated mutation rate for influenza. For higher values of *μ* (*μ* ≥ 1*e* − 4), such as those seen in HCV and HIV-1, all the tools performed poorly (i.e., both precision and recall were close to zero). For the values of *μ* seen in HIV-1 (3*e* − 5 ≤ *μ* ≤ 1*e* − 4), PredictHaplo was able to produce better results than the other tools; PredictHaplo’s precision and recall decreased with *μ* ∈ (3e−5, 1e−4) but stayed positive. It also should be noted that CliqueSNV outperformed all other tools for *μ* = 1*e* − 6, but did not produce any results for *μ* ∈ (1*e* − 5, 1*e* − 4). Such behavior looks promising and it is possible that in future releases, if run-time is increased, CliqueSNV will exceed PredictHaplo in precision and recall performance.

**Figure 3.**
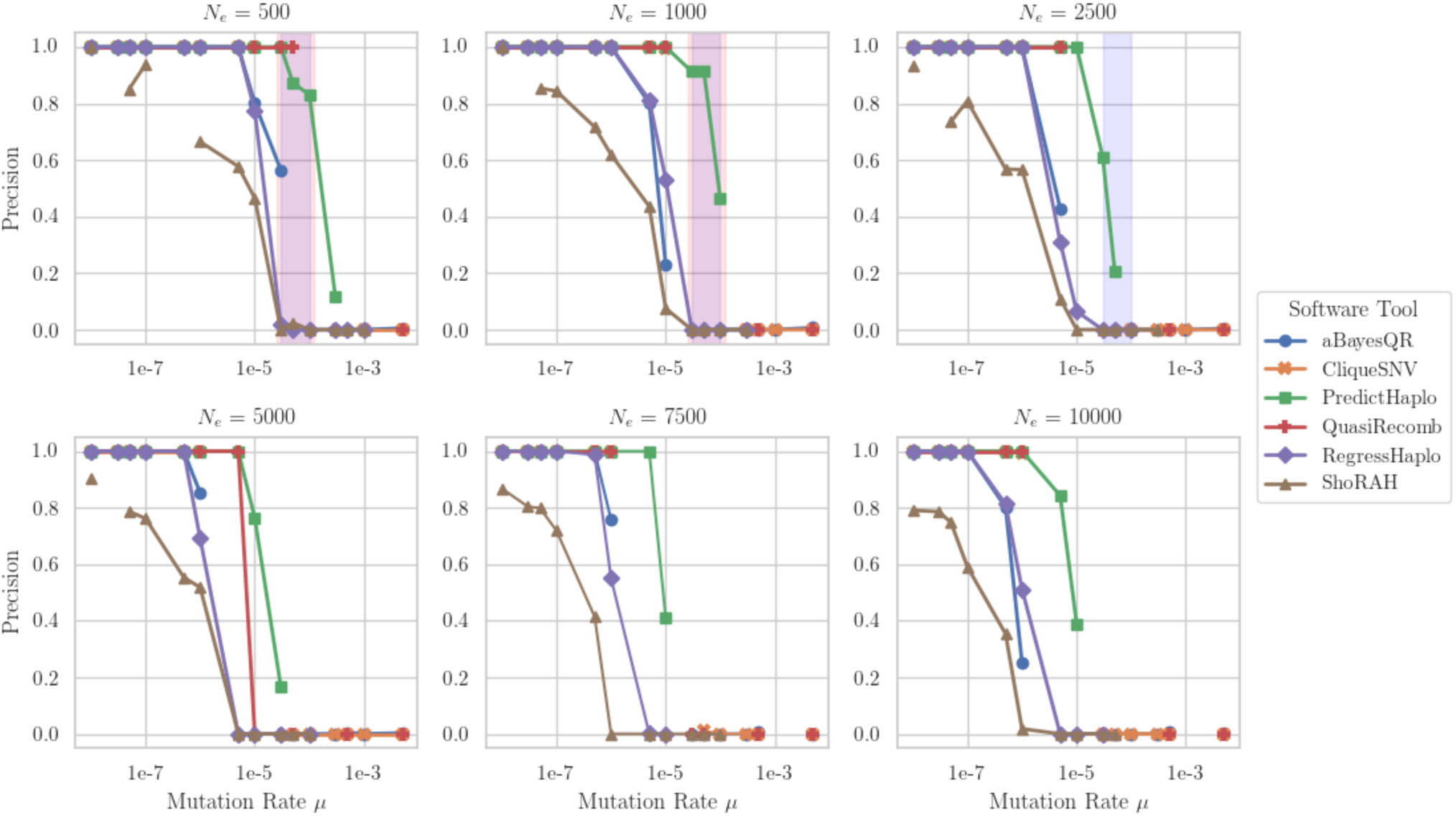
Reference-based haplotype callers: variation of precision values with mutation rate (log-scaled) for all considered *N*_*e*_. The shaded light blue and shaded light red regions correspond to HIV-1 and HCV diversity levels, respectively. For all pairs of parameters *μ* and *N*_*e*_, we report the mean estimates of precision over all valid outputs produced by each software tool for the five haplotype populations. If a tool did not produce any output for any pair of parameters, we included a gap in the corresponding plot.

**Figure 4.**
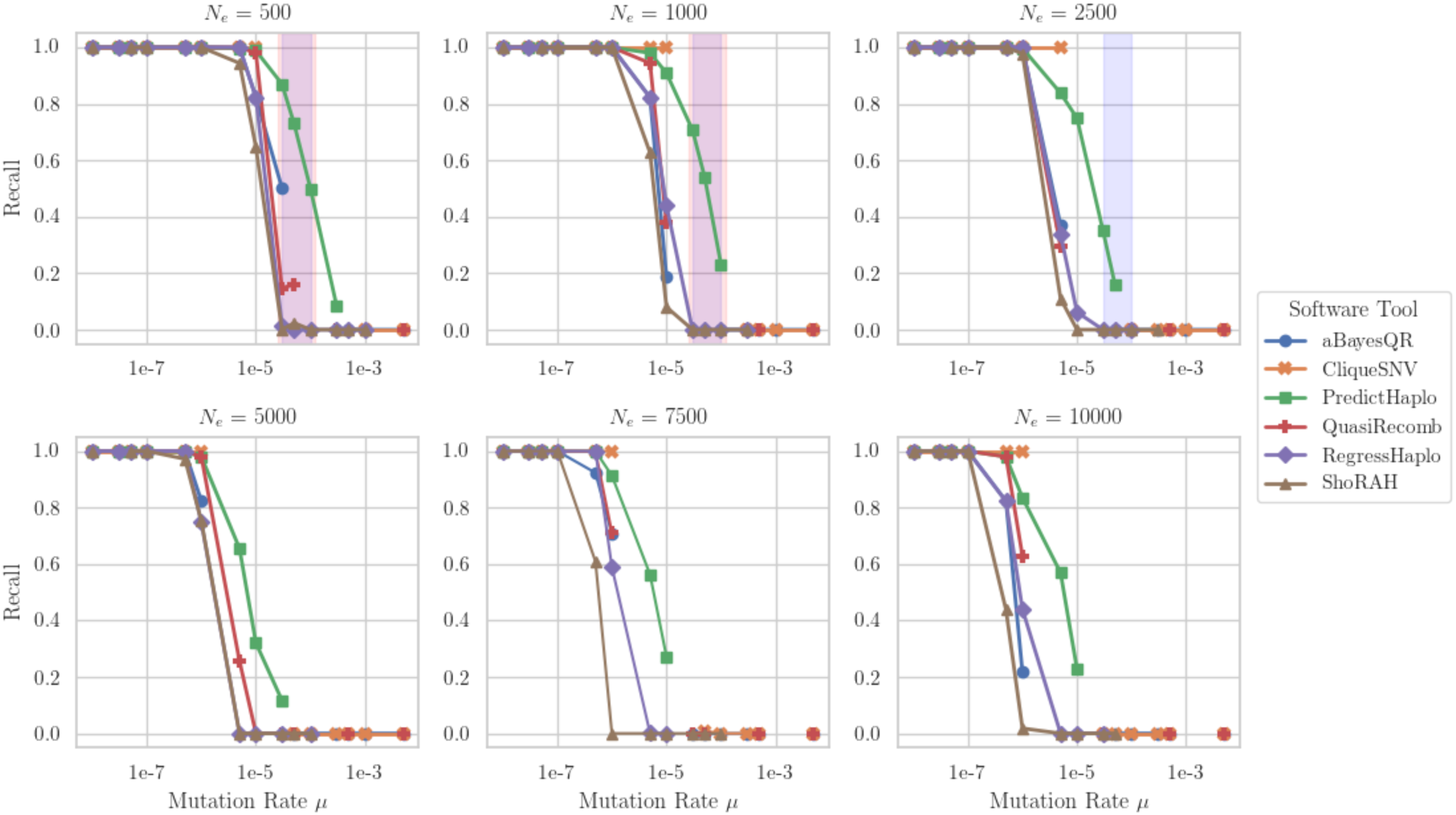
Reference-based haplotype callers: variation of recall values with mutation rate (log-scaled) for all considered *N*_*e*_. The shaded light blue and shaded light red regions correspond to HIV-1 and HCV diversity levels, respectively. For all pairs of parameters *μ* and *N*_*e*_, we report the mean estimates of recall over all valid outputs produced by each software tool for five haplotype populations. If a tool did not produce any output for some pair of parameters, we included a gap in the corresponding plot.

We calculated UniFrac distance values for the aforementioned tools (Fig. 5). The UniFrac distance further supported the previous observation that the quality of obtained results does not depend much on the effective population size (*N*_*e*_). Comparisons using the UniFrac distance also showed that all the tools, except ShoRAH, performed well if *μ* ≤ 1e – 5; the UniFrac distance between the ground truth sets of haplotypes and those predicted by the tool sets are all close to zero. With increasing *μ* values, UniFrac distances also increased. For HIV-1 mutation rates, PredictHaplo showed the best performance since it produced outputs for almost all pairs of parameters and the sets of predicted haplotypes were the closest to the correct haplotypes. Again, CliqueSNV outperformed all other methods for *μ* = 1*e* − 6, which further supports our previous observation.

**Figure 5.**
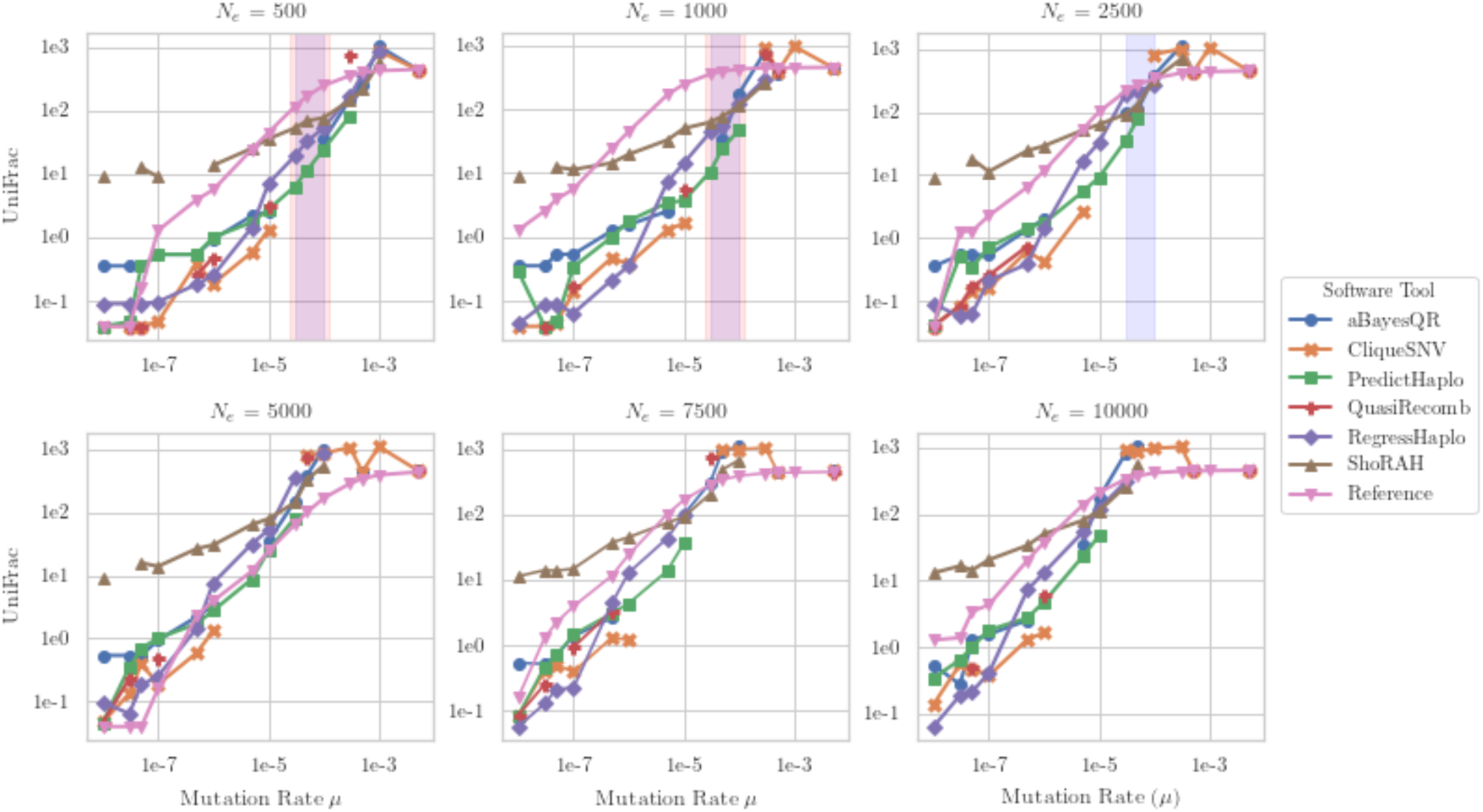
Reference-based haplotype callers: variation of UniFrac distances (EMD) with mutation rate (log-scaled) for all considered *N*_*e*_. The shaded light blue and shaded light red regions correspond to HIV-1 and HCV diversity levels, respectively. For all pairs of parameters *μ* and *N*_*e*_, we report the mean estimates of UniFrac distances over all valid outputs produced by each software tool for five haplotype populations. If a tool did not produce any output for some pair of parameters, we included a gap in the corresponding plot.

For large values of *μ* (*μ* ≥ 1*e* − 4), ShoRAH, QuasiRecomb, RegressHaplo and PredictHaplo rarely produced a valid output within the given time limit. aBayesQR and CliqueSNV produced results that were worse than or comparable to the baseline. For large values of the effective population size (*N*_*e*_ ≥ *5000*) and low values of *μ*, all the tools except ShoRAH showed better results than the baseline. However, for mutation values larger than 5*e* − 4, none of the tools made a better prediction of the set of haplotypes than just a reference. It is important to note that HCV, HIV, and influenza do not have *N*_*e*_ close to 5,000 (Bernini et al., 2011; Kim and Kim, 2016; Maldarelli et al., 2013; McCrone, 2018; McCrone et al., 2018). Most methods severely underestimated the true number of haplotypes in a population at high genetic diversity levels or overestimated it at low genetic diversity levels (Fig. 6), compared to the true number of haplotypes across the same levels of underlying genetic diversity obtained from the simulated datasets (Fig. S3). PredictHaplo, RegressHaplo, aBayesQR, and CliqueSNV underestimated haplotype numbers in the HIV intra-host diversity range (shaded in yellow). HaploClique and QuasiRecomb, on the other hand, overestimated haplotype numbers, whereas ShoRAH provided the closest estimate to the true number of haplotypes in the HIV-1 diversity range. aBayesQR and CliqueSNV did not produce results for any dataset within the HIV-1 diversity range.

**Figure 6.**
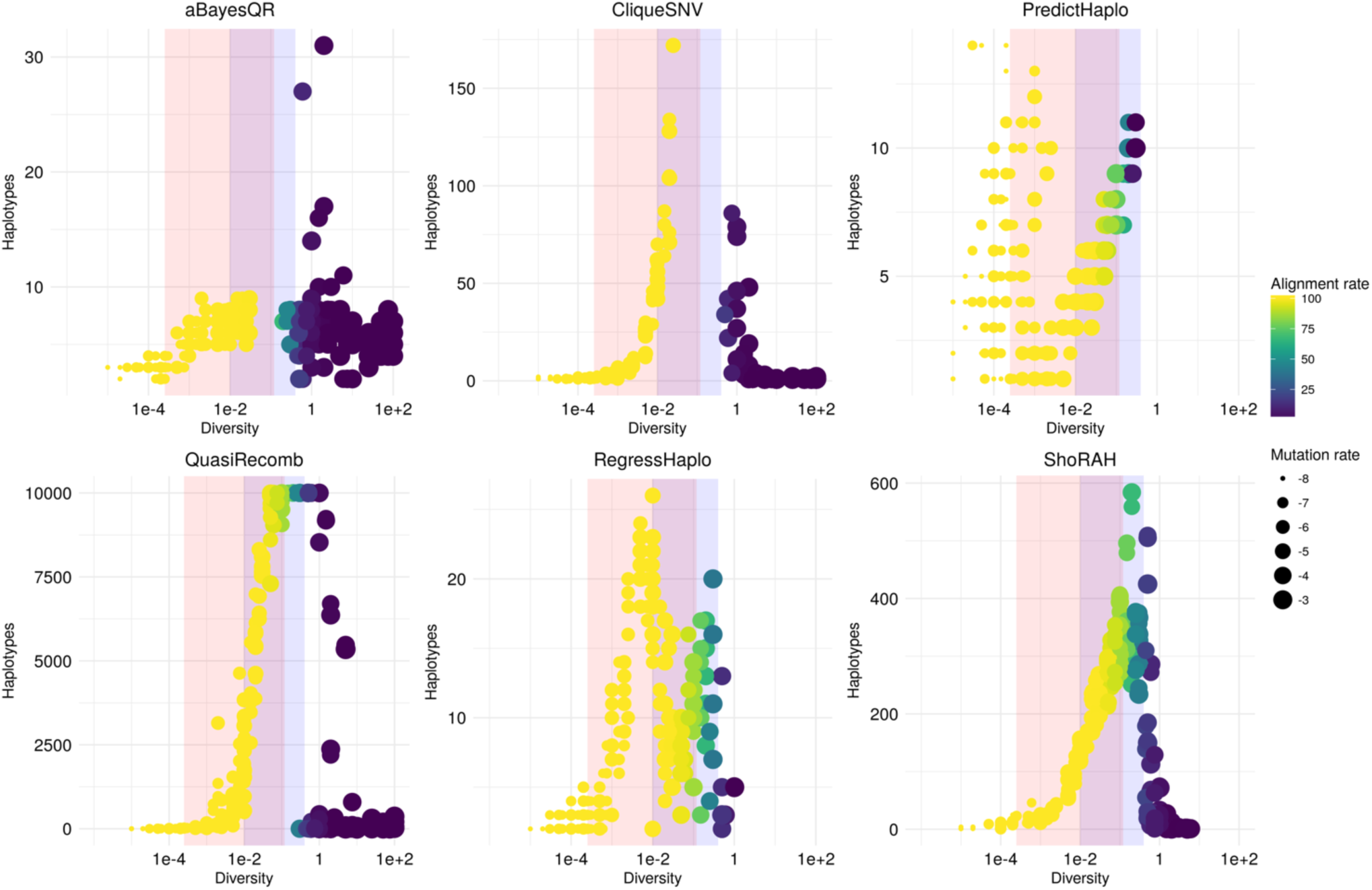
Reference-based haplotype callers: number of predicted haplotypes across levels of underlying genetic diversity. Intra-host HIV-1 and HCV diversity levels are highlighted shaded light blue and shaded light red regions, respectively. If a software tool did not complete haplotype reconstruction within the given time frame, we included a gap in the corresponding plot (see Fig. S1 for more information on dataset completions).

#### 3.2.2 De Novo Program Performances

We analyzed the behavior of two *de novo* haplotype callers: SAVAGE and PEHaplo. The output of both tools usually contained shorter contigs, so we completed the assembly using the VG-flow tool. PEHaplo itself produced valid output for all the datasets, while SAVAGE or PEHaplo with VG-flow failed to produce results for some datasets (Fig. S1). Moreover, the length of PEHaplo output haplotypes was usually closer to the ground truth haplotype length, while the SAVAGE+VG-flow produced shorter contigs (see N50 statistics plot on Fig. S4). Thus, we only further considered PEHaplo, PEHaplo + VG-flow and SAVAGE + VG-flow.

We compared the *de novo* tools using our modified versions of precision and recall (Fig. 7 and Fig. 8). VG-flow usually improved slightly the performance of PEHaplo, while PEHaplo usually performed better than SAVAGE+VG-flow. Although the quality of results of SAVAGE+VG-flow did not seem to depend on the effective population size, *N*_*e*_ played a role in the quality of obtained results by PEHaplo. For example, both precision and recall were close to zero for *μ* = 1*e* − 8 and *N*_*e*_ ∈ {500, 1000, 2500), but significantly higher for *N*_*e*_ ∈ {5000, 7500, 10000} and *μ* = 1*e* − 8. It is also important to note the behavior of recall values for the obtained results in PEHaplo; those values, in general, were close to one for small *μ* values, close to zero for *μ* values near 1*e* − 5, and stayed positive for higher *μ* values.

**Figure 7.**
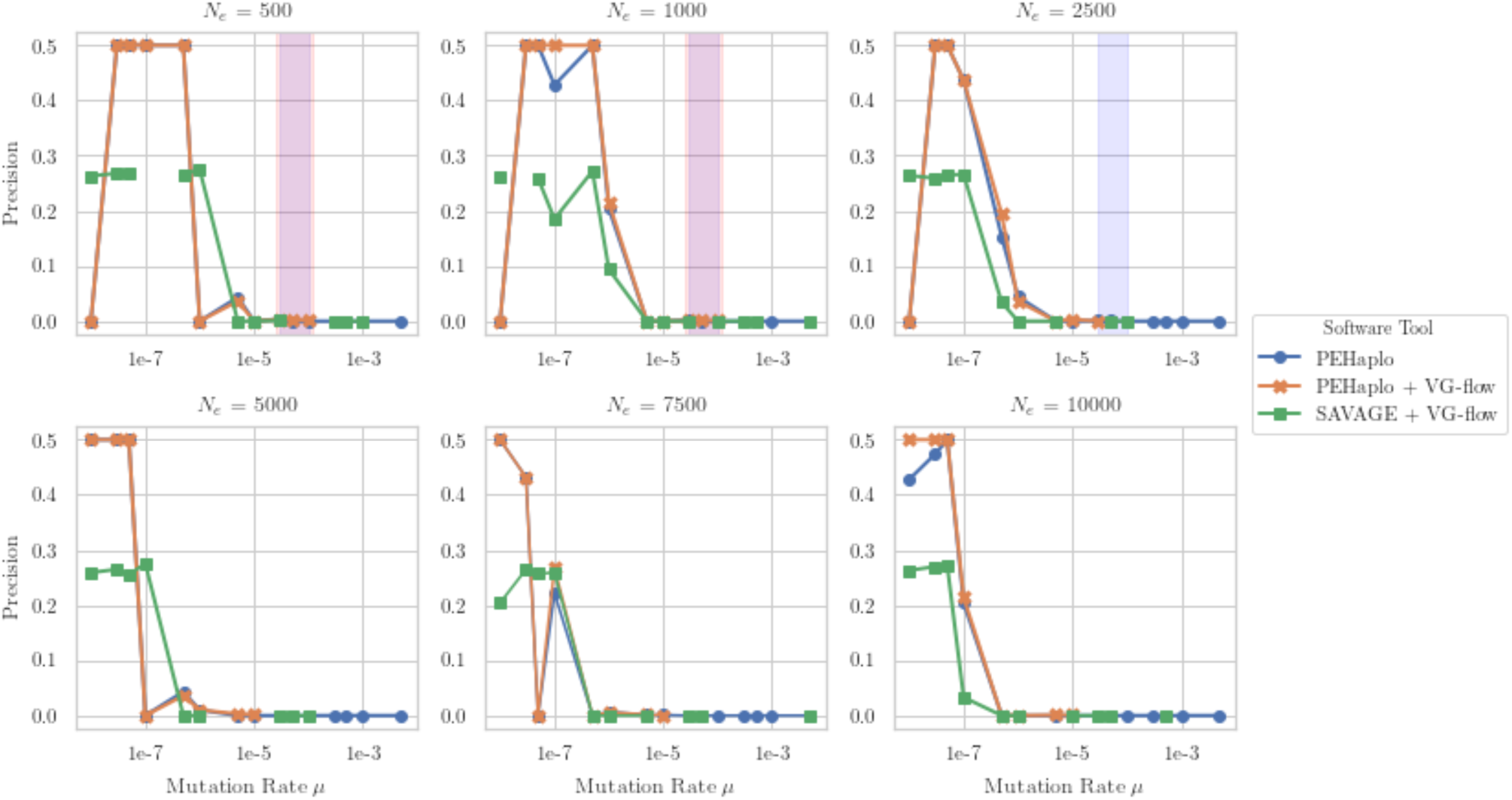
*De novo* haplotype callers: variation of precision values with mutation rate (log-scaled) for all considered *N*_*e*_. The shaded light blue and shaded light red regions correspond to HIV-1 and HCV diversity levels, respectively. For all pairs of parameters *μ* and *N*_*e*_, we report the mean estimates of precision over all valid outputs produced by each software tool for five haplotype populations. If a tool did not produce any output for some pair of parameters, we included a gap in the corresponding plot.

**Figure 8.**
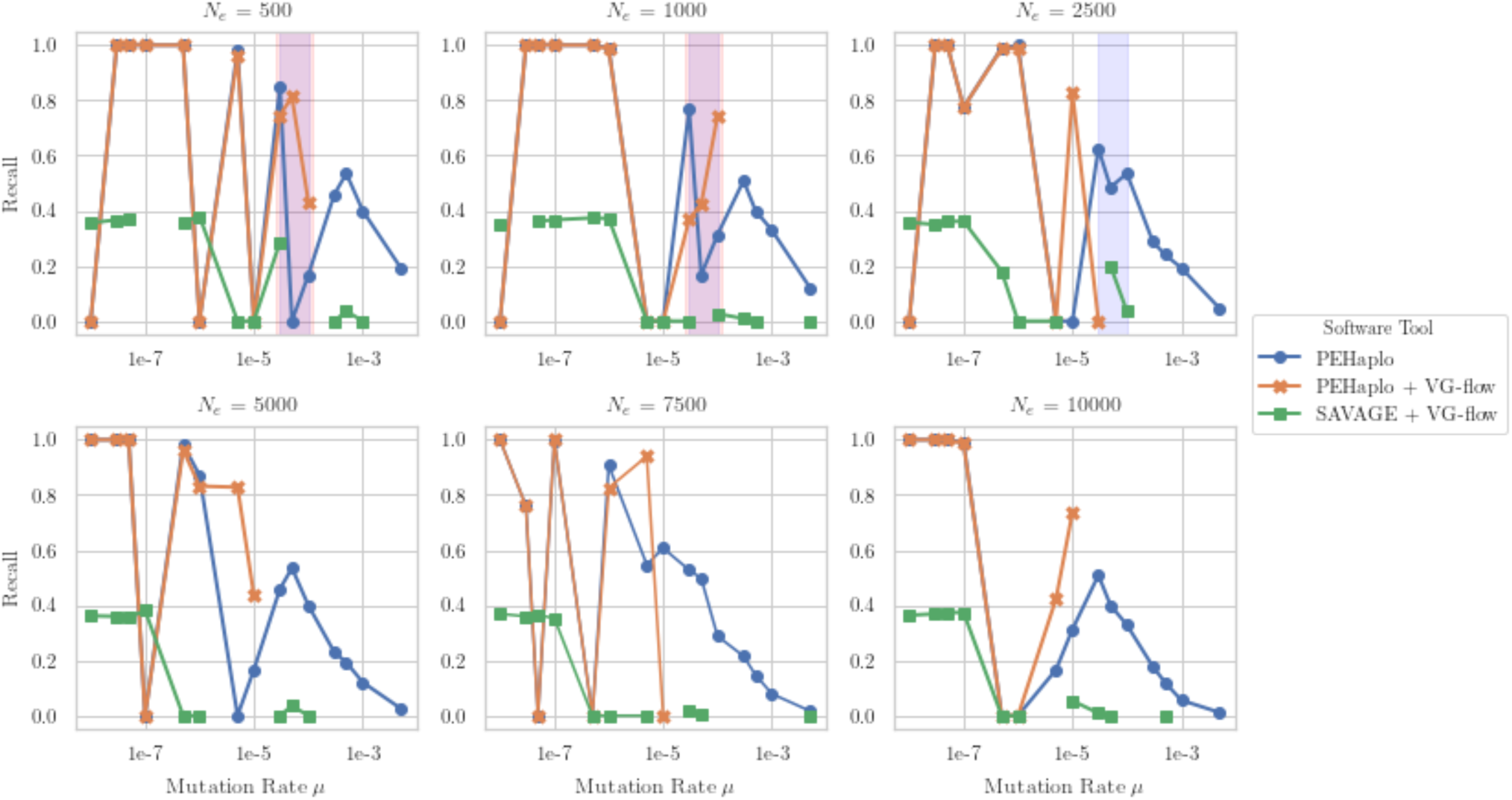
*De novo* haplotype callers: variation of recall values with the mutation rate (log-scaled) for all considered *N*_*e*_. The shaded light blue and shaded light red regions correspond to HIV-1 and HCV diversity levels, respectively. For all pairs of parameters *μ* and *N*_*e*_, we report the mean estimates of recall over all valid outputs produced by each software tool for five haplotype populations. If a tool did not produce any output for some pair of parameters, we included a gap in the corresponding plot.

*De novo* tools performed very well, in both precision and recall values, if the mutation rate was less than 1e − 6 (in contrast to *μ* ≤ 1e − 5 for *reference-based* tools). Additionally, recall values for PEHaplo when *μ* ≥ 1*e* − 4 were usually better than those seen for any *reference-based* approaches. *De novo* tools did not produce results with a positive precision for HIV-1 and HCV mutation rates. The UniFrac distance further confirmed our previous observation that VG-flow slightly improved the performance of PEHaplo (Fig. 9). Moreover, the performance of SAVAGE + VG-flow did not depend on the mutation rate or the effective population size *N*_*e*_. It is important to note that all UniFrac distance values were, in general, higher than baseline values. We also compared UniFrac distances between both categories of assemblers (Fig. S5); as we expected, *reference-based* tools largely outperformed *de novo* tools. At the same time, PEHaplo performed better than ShoRAH for some datasets. Moreover, SAVAGE + VG-flow showed the worst performance based on UniFrac distances.

**Figure 9.**
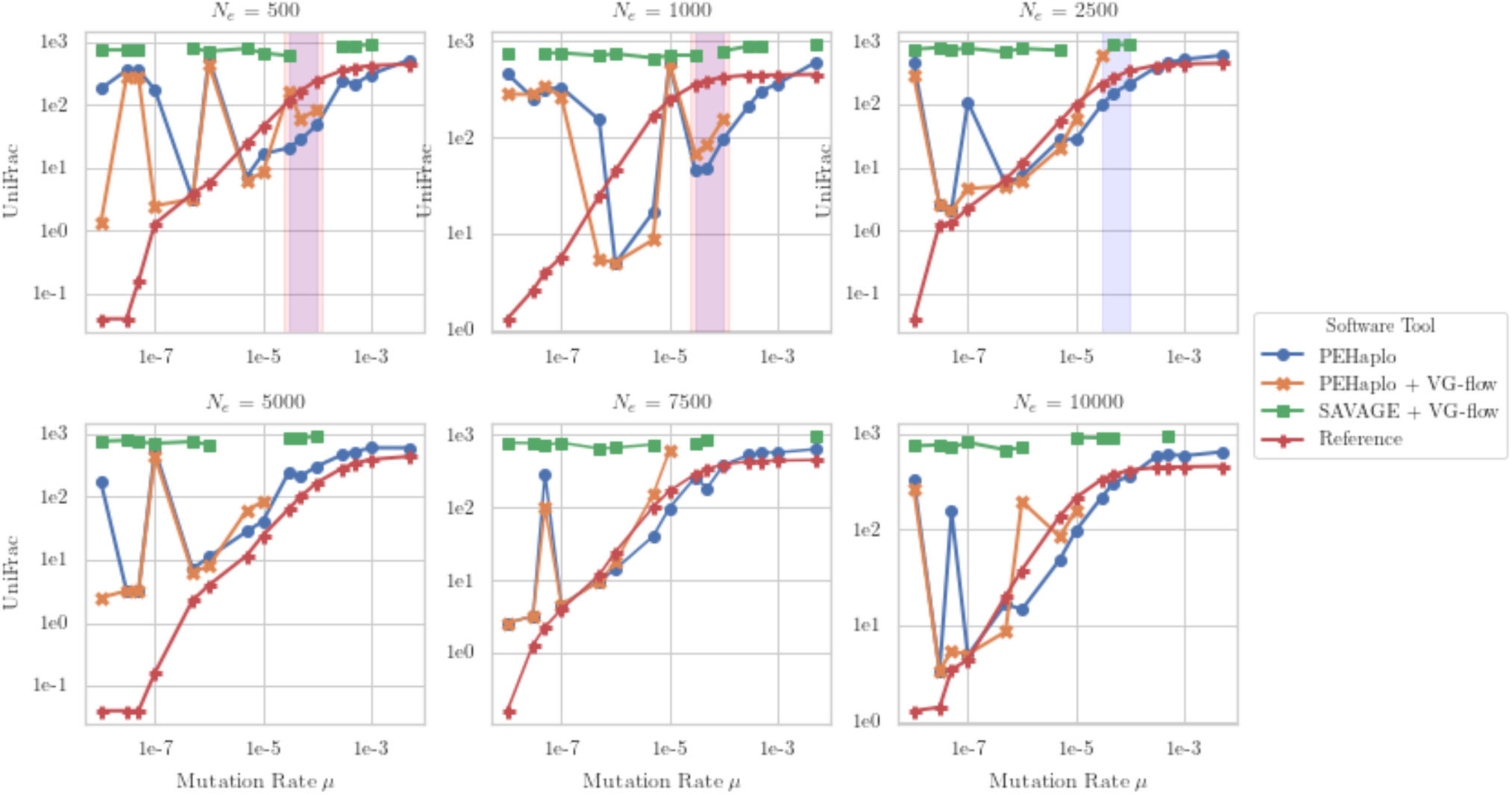
*De novo* haplotype callers: variation of UniFrac distance (EMD) with mutation rate (log-scaled) for all considered *N*_*e*_. The shaded light blue and shaded light red regions correspond to HIV-1 and HCV diversity levels, respectively. For all pairs of parameters *μ* and *N*_*e*_, we report the mean estimates of UniFrac distances over all valid outputs produced by each software tool for five haplotype populations. If a tool did not produce any output for some pair of parameters, we included a gap in the corresponding plot.

Although the *de novo* methods produced more haplotypes in the HIV-1 diversity range compared to *reference-based* methods, they all still underestimated the true number of haplotypes in a population at higher diversity levels. They also overestimated true haplotype numbers at lower genetic diversity levels (Fig. 10) compared to the true number of haplotypes from the simulated datasets (Fig. S3). When extending the contigs into scaffolds with VG-flow, the number of haplotypes reconstructed decreased considerably and remained below the number of true haplotypes estimated for the varying genetic diversity levels. PEHaplo reconstructed the lower limit of the true number of haplotypes within HIV-1 diversity levels, but like other tools, including aBayes, CliqueSNV and QuasiRecomb, PEHaplo and SAVAGE, had trouble reconstructing viral sequences at higher diversity levels.

**Figure 10.**
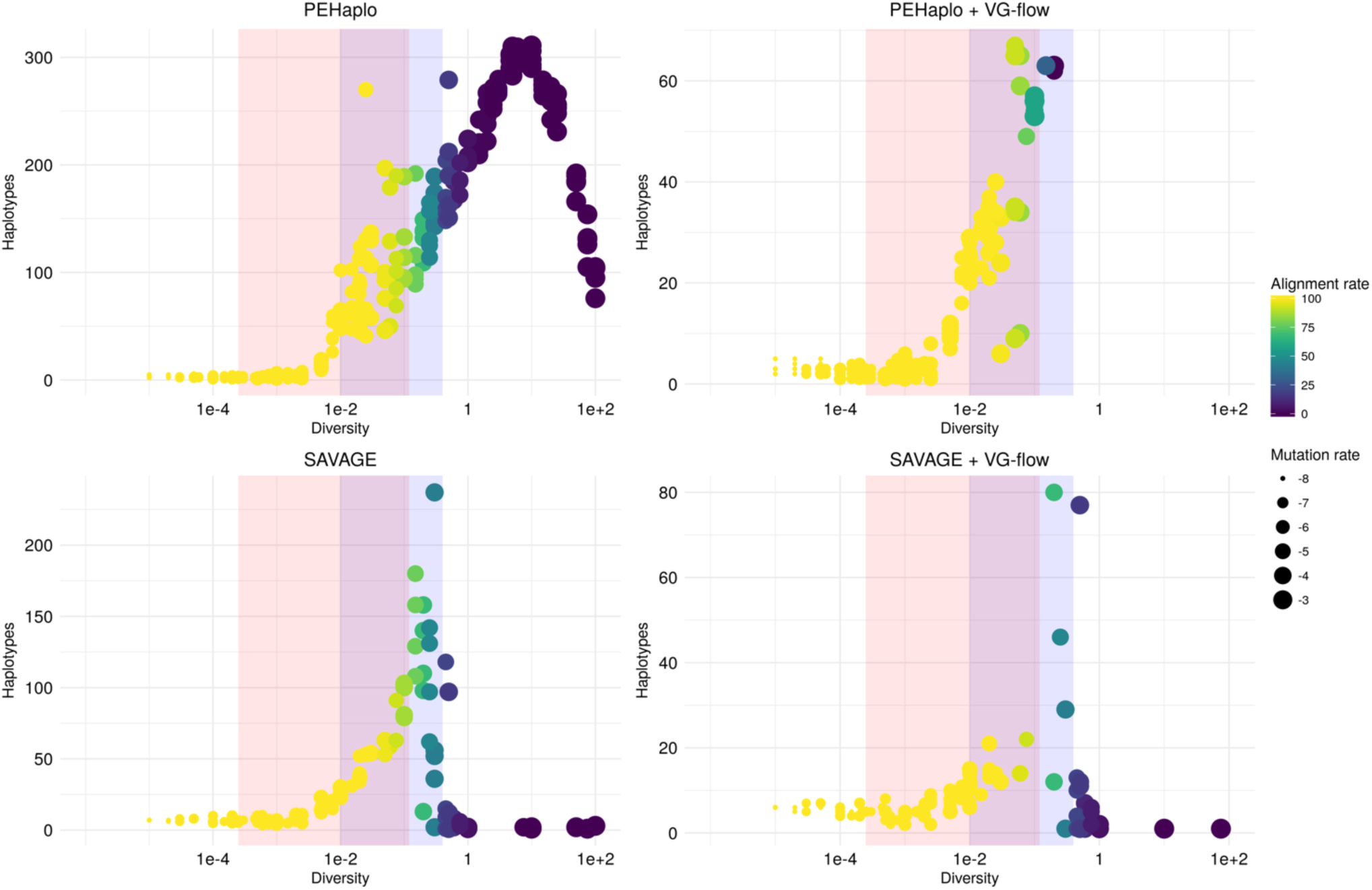
*De novo* haplotype callers: variation in number of predicted haplotypes across levels of underlying genetic diversity. Intra-host HIV-1 and HCV diversity levels are highlighted shaded light blue and shaded light red regions, respectively. If a software tool did not complete haplotype reconstruction within the given time frame, we included a gap in the corresponding plot (see Fig. S1 for more information on dataset completions).

## 4. Conclusions and Future Directions

We compared twelve of the most commonly used software tools to reconstruct haplotypes from viral NGS data. We simulated coalescent-based populations that spanned past known levels of viral diversity, including mutation rates, sample size, and effective population size. We focused our empirical comparisons on the intra-host diversity levels of fast-evolving RNA viruses such as HIV-1 because parameter value ranges are well established and a better understanding of viral dynamics is important for drug and vaccine development. Additionally, the majority of haplotype tool developers used HIV-1 to validate their own programs. In our analyses of HIV-1 intra-host diversity, we estimated between 216 and 1,185 haplotypes with a <7% frequency for a single haplotype.

Overall, *reference-based* assemblers produced more accurate haplotypes than *de novo*-based assemblers for all performance indices (precision, recall, UniFrac, and number of reconstructed haplotypes) across HIV-1 diversity levels. This performance could be attributed to the availability of high-quality reference sequences for HIV-1, HIV-2, HCV, influenza and other viruses. Furthermore, using a reference sequence reduces the computational time and power needed to reconstruct haplotypes. Reference-based assemblers likely performed better than *de novo* assemblers because of the high variation within viral populations, especially HIV-1, where the reference sequence may have provided needed guidance to orient the highly diverse NGS sequences into a haplotype sequence.

Our results show that PredictHaplo offers the best tradeoff between statistical performance and computational efficiency within HIV-1 diversity ranges. PredictHaplo was found to have the highest precision, recall, and lowest UniFrac distance values. CliqueSNV followed closely and may actually outperform PredictHaplo if more computational resources were made available. An important caveat for both these approaches, however, is that the number of true haplotypes is greatly underestimated. If it is important to identify the true number of haplotypes (as in rare haplotype discovery) approaches such as ShoRAH or PEHaplo may be more appropriate. The haplotype programs also varied greatly in terms of their ease-of-use. This variation is due to differences in coding language, program dependencies, availability of executable files, absence of comprehensive documentation and lack of example datasets. For example, SAVAGE, PEHaplo, ShoRAH can be easily installed by package managers, and CliqueSNV and QuasiRecomb are distributed as executable files. In contrast, Virus-VG and VG-flow requires installment of proprietary software, which has an academic license. Installation and usage of PredictHaplo is challenging because of the lack of description and instructions regarding the config file. While CliqueSNV is easier to install and use, there are no example datasets.

It is important to note that our study represents an initial attempt of comprehensive comparison of available haplotype reconstruction tools. For example, we focused HIV-1 diversity estimates for the polymerase gene, which is less variable than the envelope gene. Moreover, almost all developers of the aforementioned tools used the polymerase gene as a source of simulating sequencing data for assessing performance of their programs and rarely used the envelope gene for the same purposes. Given the envelope gene has a higher mutation rate and the haplotype reconstruction tools – *de novo* or *reference-based* – seem to be dependent on mutation rate, it is likely that the tools available here would not be successful in reconstructing envelope haplotypes for HIV-1 accurately. However, we chose polymerase as our gene of interest because of research focus on this gene as the target for drug resistance mutations. The same concept of lower mutation rates in conserved genes and higher mutation in less conserved genes can be seen in other fast-evolving viruses. For example, in HCV the core protein is more conserved compared to the E1/E2 region. Thus, users should target methods for haplotype calling that best match the mutation rate of their target gene.

Another limitation of our study is coverage. It is well-known that coverage plays a crucial role in all algorithms for distinguishing between errors and rare sequence variants. We chose 100x coverage because it represents a reasonable amount of data that can be obtained without intensive labor or money consuming procedures. Contrary to our simulations, the developers of haplotype reconstruction tools usually test their methods on datasets with higher coverage than ours. For example, the famous golden-standard benchmark HIV dataset (*labmix* dataset (Di Giallonardo et al., 2014)) on which all tools have been tested by developers, consisted of an average of 20,000x coverage. Thus, our study represents an attempt to measure the performance of the haplotype reconstruction tools on datasets that are more likely to be seen and produced in laboratories. Moreover, according to our results for higher mutation rates, many tools did not produce any results within the time limit. Considering that higher coverage implies a larger amount of data and thus requiring more computational time to process these data, it is expected that the tools available here would require extensive computational resources.

We also considered error-free and recombination-free data in our study. Only a few tools explicitly took into account the presence of errors or recombination in their models (e.g., only QuasiRecomb explicitly assumes the presence of recombination events). By not simulating recombination and sequencing errors, we removed nuisance parameters that would impact haplotype reconstruction. Moreover, since almost all tools have been tested on ultra-deep data, our comparison study by error-free data is giving an advantage to these methods by removing errors in sequence data (i.e., one does not need deep coverage to distinguish between rare variants and errors). Furthermore, Zanini et al. (2015) found evidence that recombination likely interrupts haplotypes, specifically in HIV-1, every 100-200bp, so, the concept of haplotypes in HIV-1, and maybe other fast-evolving viruses with high recombination rates, may not exist or be feasible to study with frequent recombination events. Together these facts imply that the performance of the aforementioned tools would be even worse than observed here.

Overall, results and limitations of our study indicate the importance of creating broad and diverse golden-standard datasets that must include several different genes, diverse parameters of mutation rates and effective population sizes, different average coverages, presence or absence of recombination events or/and error prone data. Moreover, future simulation studies should address error-prone data using haplotype callers that can handle sequencing errors and investigate the effect of recombination and average coverage on the reconstruction of haplotype. In addition to simulation studies, some theoretical work similar to DNA sequencing theory should be done for laying analytical foundations for determining coverage depending on the mutation rate, effective population size, error-rate of a sequencing technology, and so on. Finally, there are still a lot of opportunities for developing new haplotype callers that can process a wide range of data with different mutation rates, average coverage, and presence or absence of recombination events. Moreover, since the reconstructed haplotypes are often used for reconstructing phylogeny, the future tools may also consider the problem of reconstructing haplotype sequences together with their phylogeny. Considering the possibility that the reconstructing haplotype sequences from short-read sequencing technologies may represent an intractable problem, focusing on reconstructing haplotype phylogeny directly from short-reads may lead to better results after all. In addition to mentioned future directions, the advances and price-decreasing of long-read sequencing technologies (e.g., Nanopore, PacBio, 10X Genomics) poses a whole new set of challenges for haplotype reconstructions including the development of new sequencing protocols and haplotype reconstruction tools. This new technology has the power to sequence long amplicons or even entire viral genomes in a single pass, i.e., no need to assemble sequencing reads. However, this type of data requires development of new methods that can distinguish between rare variants and sequencing errors. Therefore, the application of long-read sequencing technology may be more beneficial for studying global, or entire genome, haplotypes.

## Abbreviations

NGS: Next generation sequencing
HIV: Human immunodeficiency virus
HCV: Hepatitis C virus
HPV: Human papillomavirus

## Acknowledgements

The authors would like to thank Adam Kai Leung Wong for his invaluable help installing and troubleshooting different software packages in the Colonial One high performance computing cluster at The George Washington University.

## Authors’ contributions

KMG, MLB, and KAC developed project idea. KAC and MP-L acquired funding for this study. KMG and MLB completed the simulation study. AE and DN ran all the haplotype reconstruction tools. AE, PA, and NA performed the statistical analysis of the results. PA and NA supervised AE and DN. PA, NA and KMG developed initial manuscript draft. All authors read and reviewed final draft of manuscript.

## Ethics approval and consent to participate

Not applicable.

## Consent for publication

Not applicable.

## Appendix

**Figure S1.**
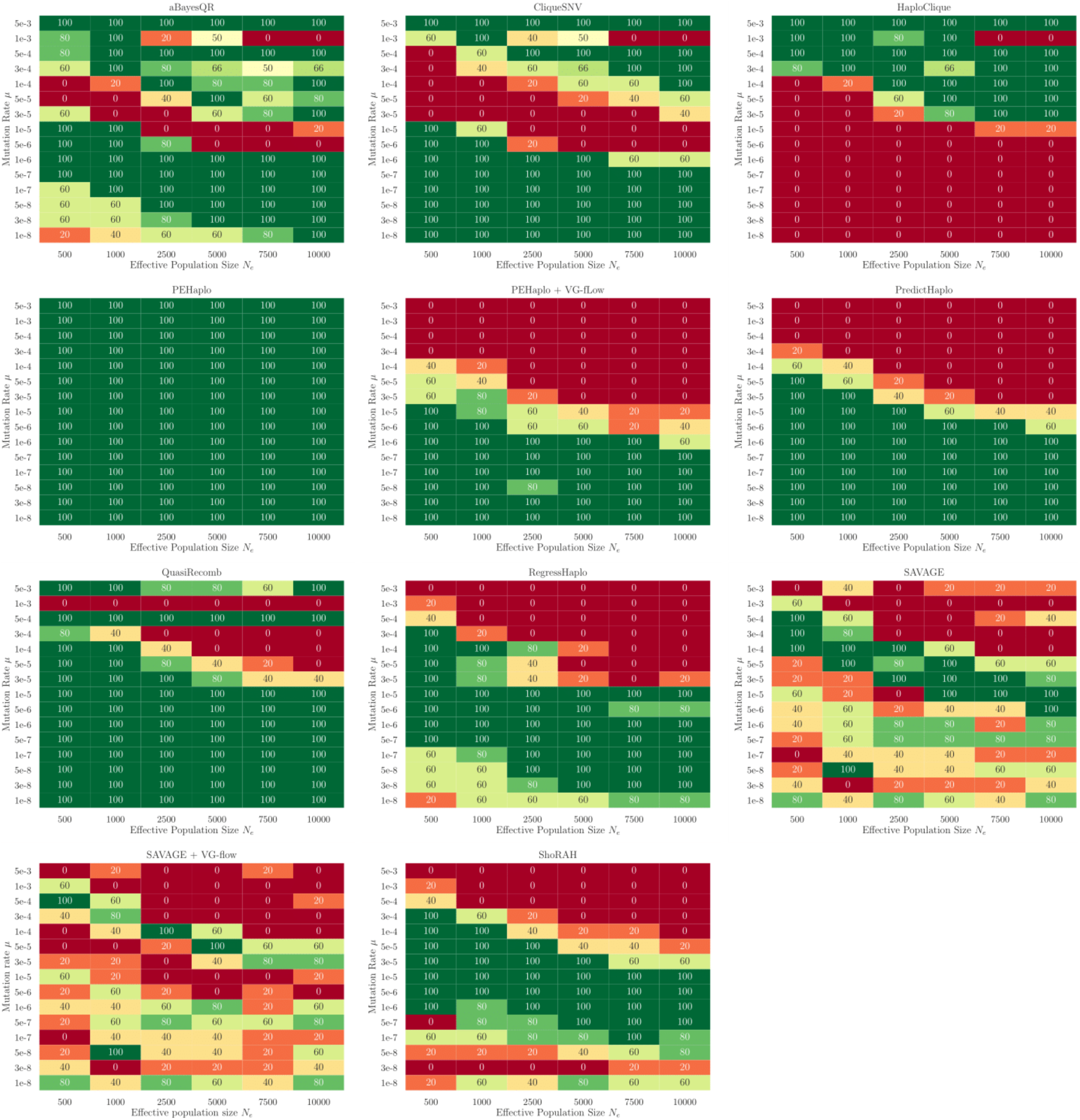
Plots showing percentage of datasets analyzed by a tool within our time limit. From green to red, full completion of a dataset (green) to no completion (red) is indicated for each set.

**Figure S2.**
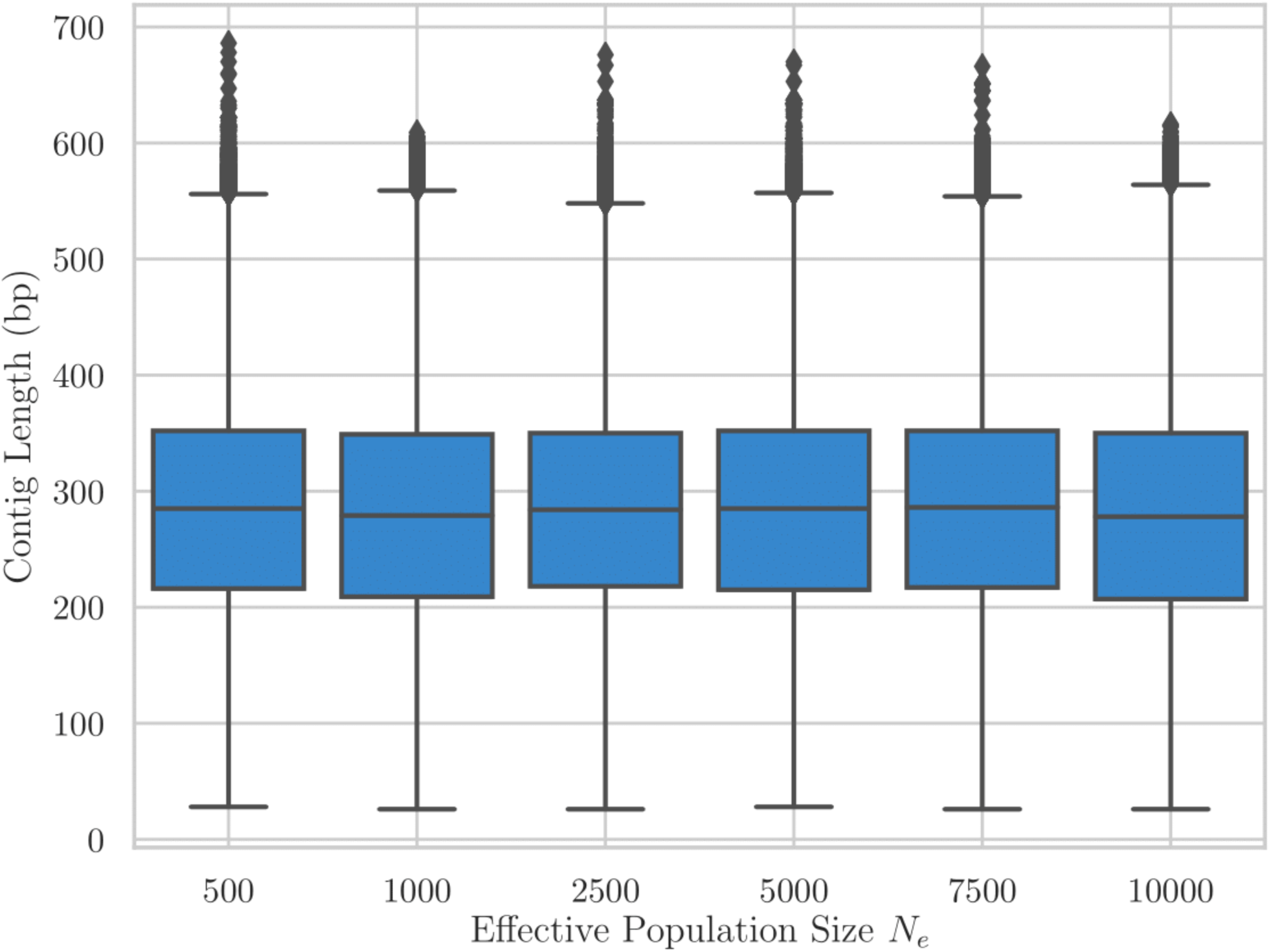
The length distribution of haplotypes obtained by HaploClique for each effective population size.

**Figure S3.**
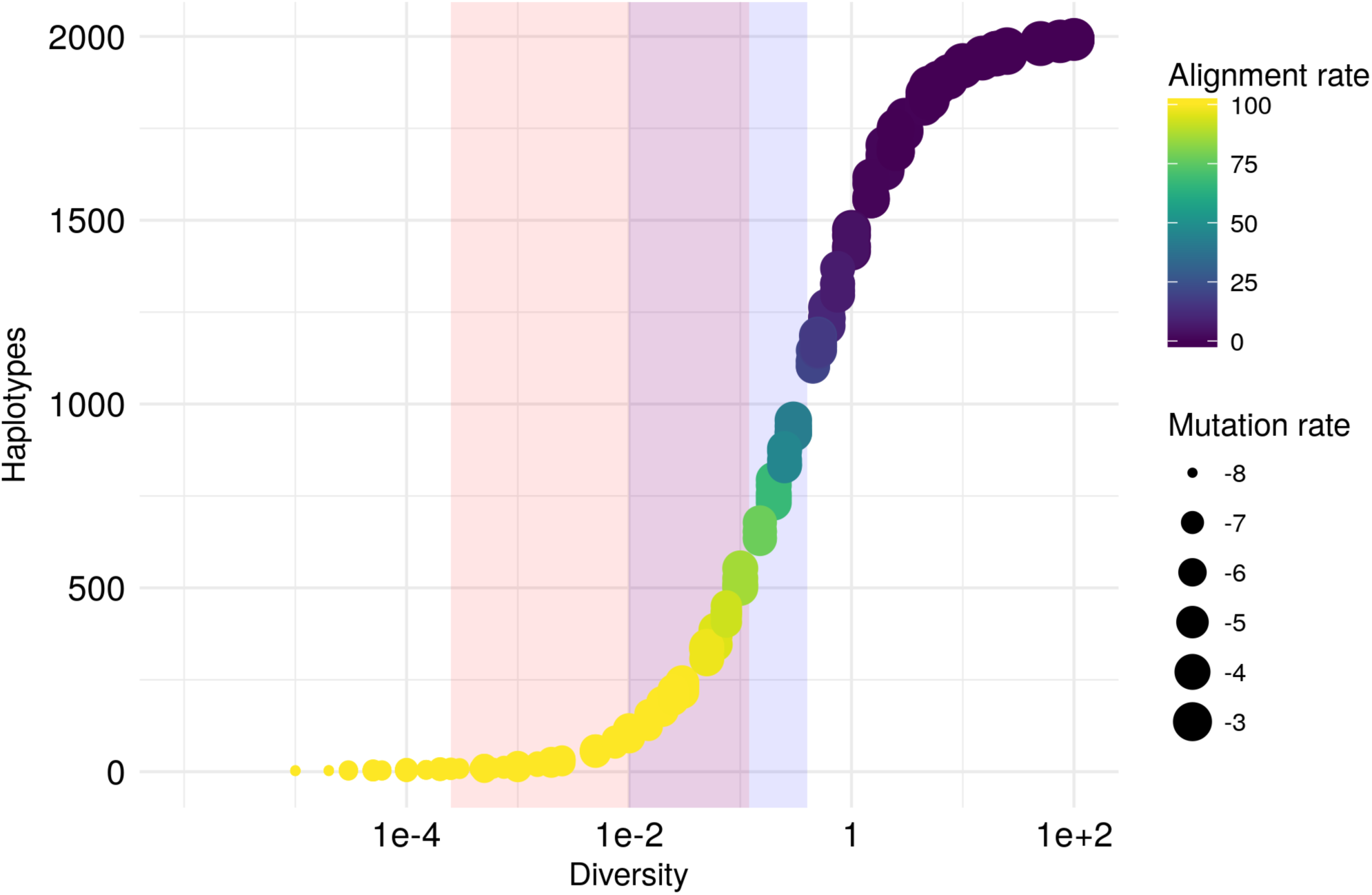
Number of true haplotypes estimated from the coalescent-based simulated data across levels of underlying intra-patient genetic diversity. Intra-host HIV-1 and HCV diversity levels are highlighted shaded light blue and shaded light red regions, respectively.

**Figure S4.**
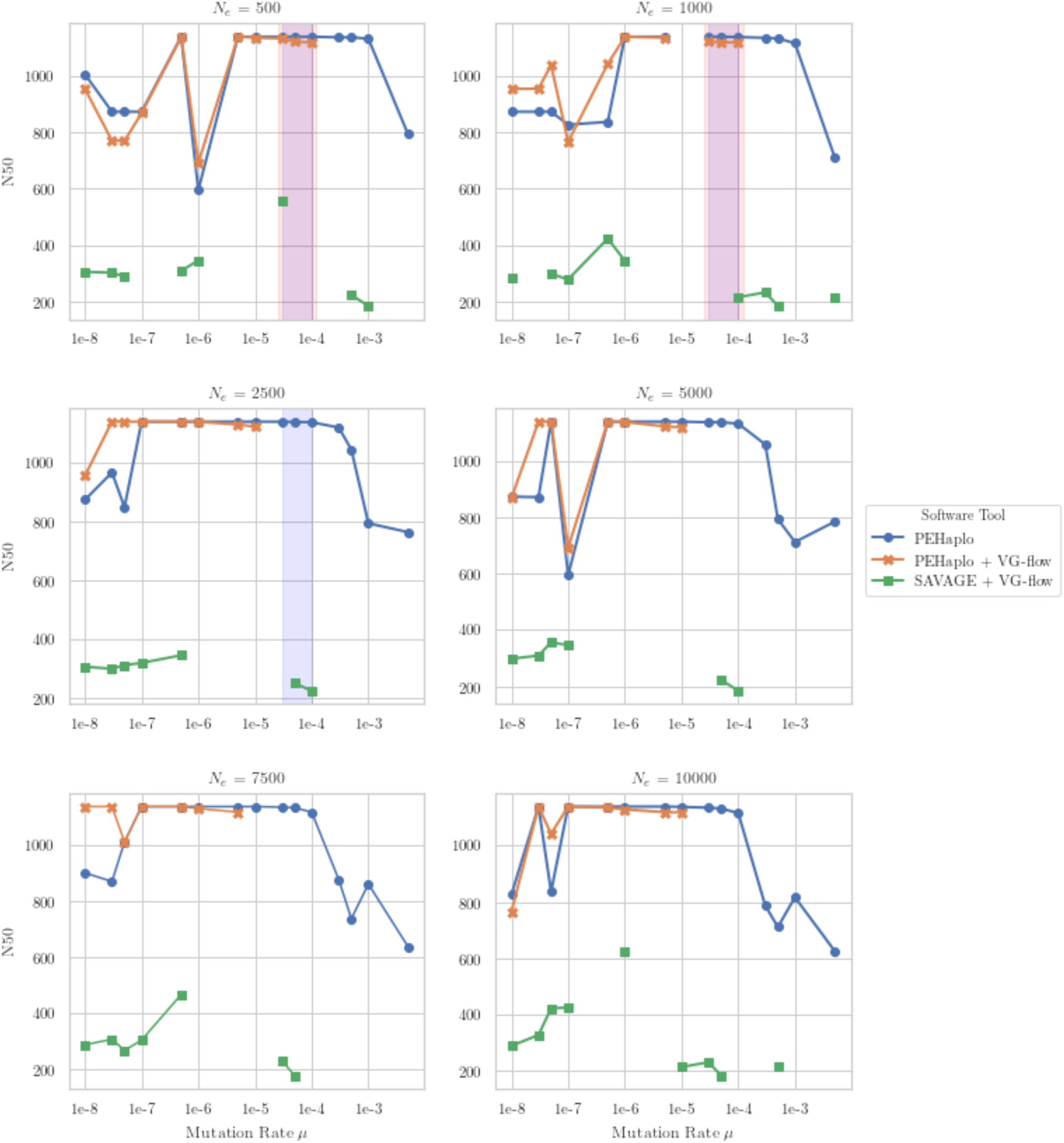
Reference-based and *de novo* haplotype callers: variation of recall values with mutation rate (log-scaled) for all considered *N*_*e*_. The shaded light blue and red regions correspond to HIV-1 and HCV diversity levels, respectively. For all pairs of parameters *μ* and *N*_*e*_, we report the mean estimates of recall over all valid outputs produced by each software tool for five haplotype populations. If a tool did not produce any output for some pair of parameters, we included a gap in the corresponding plot.

**Figure S5.**
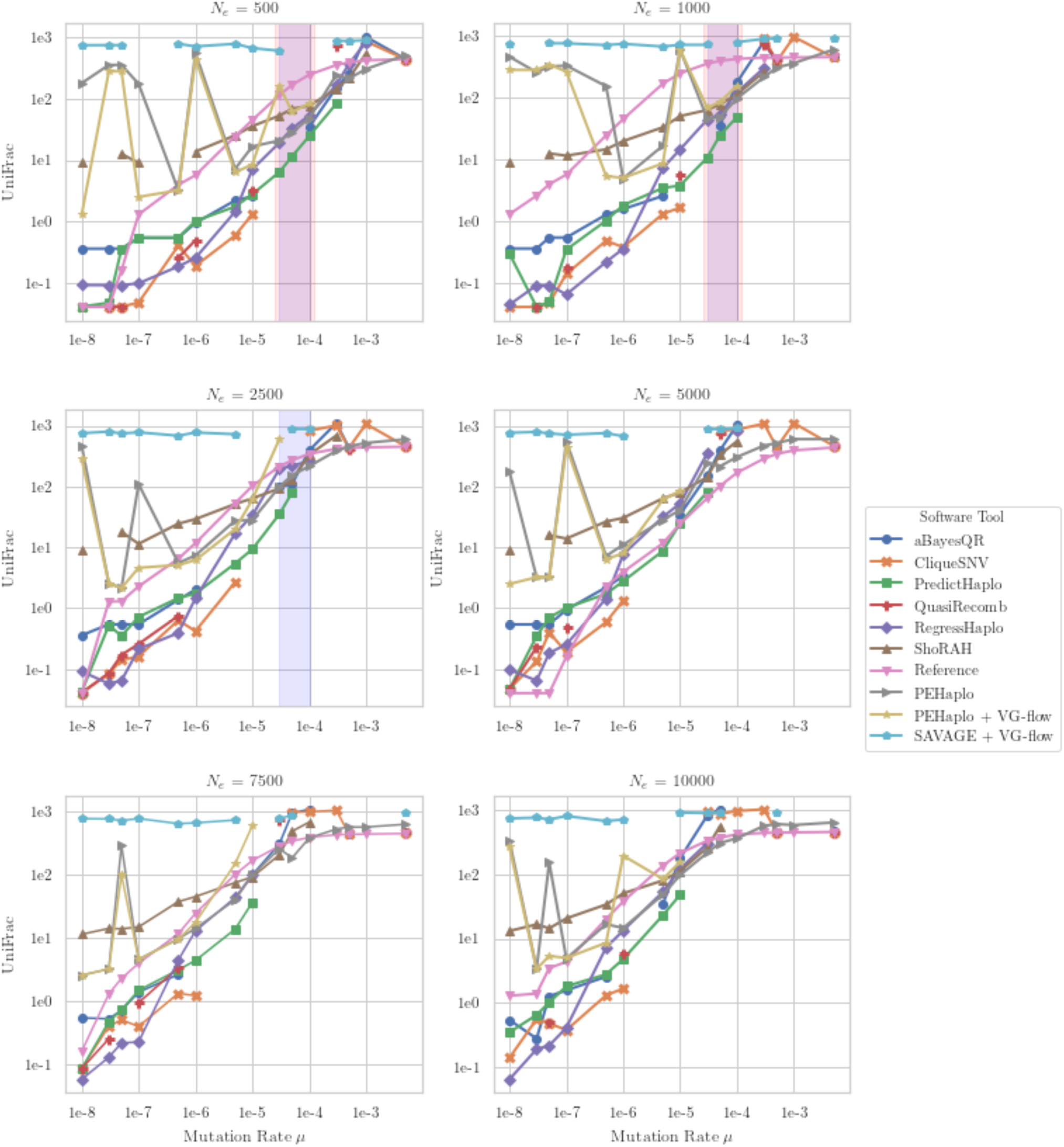
Reference-based and *de novo* haplotype callers: variation of UniFrac distances (EMD) with mutation rate (log-scaled) for all considered *N*_*e*_. The shaded light blue and red regions correspond to HIV-1 and HCV diversity levels, respectively. For all pairs of parameters *μ* and *N*_*e*_, we report the mean estimates of UniFrac distances over all valid outputs produced by each software tool for five haplotype populations. If a tool did not produce any output for some pair of parameters, we included a gap in the corresponding plot. The values for *de novo* tools are usually higher than for *reference-based* tools.

